# Benchmarking *de novo* assembly methods on metagenomic sequencing data

**DOI:** 10.1101/2022.05.22.493002

**Authors:** Zhenmiao Zhang, Chao Yang, Xiaodong Fang, Lu Zhang

## Abstract

Metagenome assembly is an efficient approach to deciphering the “microbial dark matter” in the microbiota based on metagenomic sequencing, due to the technical challenges involved in isolating and culturing all microbes in vitro. Although short-read sequencing has been widely used for metagenome assembly, linked- and long-read sequencing have shown their advancements by providing long-range DNA connectedness in assembly. Many metagenome assembly tools use dedicated algorithms to simplify the assembly graphs and resolve the repetitive sequences in microbial genomes. However, there remains no comprehensive evaluation of the pros and cons of various metagenomic sequencing technologies in metagenome assembly, and there is a lack of practical guidance on selecting the appropriate metagenome assembly tools. Therefore, this paper presents a comprehensive benchmark of 15 de novo assembly tools applied to 32 metagenomic sequencing datasets obtained from simulation, mock communities, or human stool samples. These datasets were generated using mainstream sequencing platforms, such as Illumina and BGISEQ short-read sequencing, 10x Genomics linked-read sequencing, and PacBio and Oxford Nanopore long-read sequencing. The assembly tools were extensively evaluated against many criteria, which revealed that compared with the other sequencing technologies, long-read assemblers generated the highest contig continuity but failed to reveal some medium- and high-quality metagenome-assembled genomes (MAGs). In addition, hybrid assemblers using both short- and long-read sequencing were promising tools to both improve contig continuity and increase the number of near-complete MAGs. This paper also discussed the running time and peak memory consumption of these tools and provided practical guidance on selecting them.

## Introduction

During long-term and complex genetic evolution, human and non-human animals have formed an ecosystem of symbiotic relationships with diverse microbes. Identifying these microbes and their genome sequences is essential for revealing their interactions with the hosts and providing rich information about human health and diseases^1–3^. The traditional identification strategy isolates and cultures the target microbes in vitro, and then each microbe’s genome is sequenced^4^. However, most microbes in some specific conditions, e.g. human gastrointestinal tract^5,6^, cannot be cultured in the laboratory, which prevents the complete deciphering of the microbial community^7^. Alternatively, metagenomic sequencing enables the efficient, direct sequencing of a mixture of microbial DNAs without the need for microbial isolation, and thus facilitates the deciphering of microbial genomes with diverse characteristics. The metagenomic sequencing data is then subjected to metagenome assembly, which aims to reconstruct microbial genomes by concatenating the sequencing reads into long genome fragments (contigs). Metagenome assembly has been proven to be an effective strategy to explore microbial genomes and their biological functions from metagenomic sequencing data^8^.

Short-read sequencing is the most prevalent technology adopted in metagenomic studies. Many tools have been developed to assemble short-reads from the microbial genomes with imbalanced coverage and to distinguish the origins of reads based on their sequence characteristics. For example, Meta-IDBA^9^ partitions *de Bruijn* graphs into isolated graph components based on their sequence similarities, with each graph component representing a unique species, and then uses each component’s consensus sequences to form a draft genome. IDBA-UD^10^ resolves short repeats from low-depth regions by local assembly using the paired-end constraint of short-reads. metaSPAdes^11^ extends SPAdes^12^ by incorporating a series of graph simplification strategies to separate strains with similar sequences, and applies ExSPAnder^13^ to detangle the repetitive sequences. MEGAHIT^14^ constructs succinct *de Bruijn* graphs using *k*-mers (subsequences of length *k*) with various *k* to fill the gaps in low-depth regions and resolve genomic repeats. Currently, the most widely used commercial short-read sequencing platforms are designed by Illumina (e.g., HiSeq, NextSeq, and MiSeq) and BGI (e.g. BGISEQ-500, MGISEQ-200 and MGISEQ-2000), which provide read lengths between 100 bp and 300 bp^15^. The following three tasks in metagenome assembly cannot be easily achieved using short read lengths, as long-range genomic connectedness cannot be determined: (1) the detection of horizontal gene transfers and transposon mobilization between species; (2) the deconvolution of duplicate and conserved sequences in microbial genomes; and (3) the generation of high-quality draft genomes for low-abundance species. As such, the advanced sequencing technologies, such as linked- and long-read sequencing, combine long-range DNA connectedness with sequencing reads and are advantageous for resolving complex genomic regions and generating more complete draft genomes than short-read sequencing.

Linked-read sequencing technology attaches short-reads with the same barcodes if they are derived from the same long DNA fragment; therefore, a barcode may include several “virtual” long-reads. Several linked-read sequencing platforms have been developed, such as Illumina TrueSeq Synthetic Long-Reads (SLR), LoopSeq, 10x Genomics (10x), single-tube Long Fragment Read (stLFR), and Transposase Enzyme-Linked Long-read Sequencing (TELL-Seq) and some of them have been used effectively for metagenome assembly, especially for the linked-reads from 10x platform. For example, Zlitni et al. explored temporal strain-level variants in stool samples from a patient during his 2-month hematopoietic cell transplant treatment, by assembling 10x metagenomic sequencing data^16^. Roodgar et al. investigated the responses of human gut microbiome during antibiotic treatment using longitudinal 10x linked-read sequencing and assembly^17^. Two assemblers have been developed for 10x linked-read sequencing data. cloudSPAdes^18^ models linked-read metagenome assembly as a shortest cloud superstring problem and leverages the reconstructed fragments to simplify the assembly graph. Bishara et al. developed Athena^19^ to improve metagenome assembly by considering co-barcoded reads for local assembly and demonstrated that 10x linked-reads outperformed Illumina short-reads and SLR in the assembly of metagenomic data from human stool samples.

Long-read sequencing technology has recently received increasing attention. Single-molecule long-read sequencing platforms, such as the PacBio Single-Molecule Continuous Long Read sequencing (PacBio CLR), the PacBio highly accurate long-read sequencing (PacBio HiFi), and the Oxford Nanopore Technologies sequencing (ONT), have been used to assemble the genomes of many isolated microbes. Chin et al. implemented an assembly tool based on a hierarchical and greedy strategy and demonstrated its ability to assemble the complete genome of Escherichia coli K-12 from the deeply sequenced PacBio CLR data^20^. A follow-up study showed that Nanocorrect and Nanopolish exhibited comparable performance in E. coli genome assembly from polished ONT long-reads^21^. Accordingly, as mentioned, long-read sequencing has recently been applied to metagenome assembly. Tsai et al. used PacBio CLR long-reads to assemble the metagenomic sequencing data of human skin, and thereby identified a previously uncharacterized Corynebacterium simulans^22^. Another recent study investigated 2,267 bacteria and archaea and found that a majority of their genomes could be assembled perfectly using PacBio CLR long-reads^23^. Many software have been developed for long-read metagenome assembly. metaFlye^24^ extends Flye^25^ to deal with uneven bacterial composition and intra-species heterogeneity by leveraging unique paths in repeat graphs. Canu^26^ improves Celera^27,28^ to deal with noisy long-reads by using multiple rounds of reads error-correction. Moss et al. improved the long-read sequencing protocol of DNA extraction and developed Lathe to optimize metagenome assembly on ONT data^29^. Several other state-of-the-art long-read assemblers have been released recently, including MECAT2^30^, NECAT^31^, Shasta^32^, and wtdbg2^33^, but few of them have been tested on metagenome assembly. Nevertheless, there are some limitations to long-read sequencing that restrict its practical application in metagenome assembly. First, the high base-error rate of long-read sequencing makes it challenging to distinguish strains and substrains with similar sequence characteristics. Second, the high cost of long-read sequencing prevents its wide application in large cohort studies.

Hybrid assembly is an alternative strategy that combines the strengths of both short-reads and long-reads. For example, the hybrid assembly tool DBG2OLC^34^ aligns the contigs assembled from short-reads to long-reads, in which the identifiers of aligned contigs are used to represent each long-read. The overlap-layout-consensus approach is then performed on the long-reads by matching their contig identifiers. metaFlye has a hybrid assembly module (“--subassemblies”) that considers the contigs assembled from both short- and long-reads as high-quality long fragments. OPERA-LG^35^ links and orientates the contigs from short-read assembly by paired-end constraint and long-reads support. OPERA-MS^36^ constructs a scaffold graph by linking the contigs assembled from short-reads if they are supported by long-reads. The contigs are grouped into clusters based on graph topology and read depths, with each cluster representing one species. Then, by leveraging reference genomes, OPERA-MS recognizes contigs derived from the same microbial strain, and OPERA-LG assembles these contigs.

Currently, the advantages and limitations of existing sequencing platforms and corresponding assembly tools remain unclear, and there is an urgent need for practical guidelines on how to select the best platform and tool for specific purposes. Sczyrba et al. evaluated several short-read assemblers using the Critical Assessment of Metagenome Interpretation (CAMI)^37^; Meyer et al. added long-read assemblers using CAMI II datasets in a follow-up study^38^. Latorre-Pérez et al. benchmarked long-read assemblers using ONT data from ZymoBIOMICSTM Microbial Communities^39^. The assembly tools they evaluated were incomplete and one-sided, and these studies used datasets from simulation or mock communities, which could not fully represent real microbial communities.

In this study, we benchmarked 15 state-of-the-art tools to generate short-read, linked-read, long-read, and hybrid assemblies using metagenomic sequencing datasets from simulation, mock communities, and human stool samples (Fig. 1). This benchmark involved comparing the basic contig statistics, including total assembly length (*AL* for human stool datasets), genome fraction (*GF* for simulation and mock datasets), and contig N50, NA50, and normalized NGA50 (**Methods**). We also evaluated the metagenome-assembled genomes (MAGs) after contig binning with respect to their continuities (MAG N50), qualities (**Methods**; *#MQ*: the number of medium-quality MAGs; *#HQ*: the number of high-quality MAGs; *#NC*: the number of near-complete MAGs), and annotations (the number of MAGs that can be annotated into species). Our results showed that the short-read assemblers generated the lowest contig continuity and *#NC*. MEGAHIT outperformed metaSPAdes on deeper sequenced datasets (>100x), and metaSPAdes obtained better results than MEGAHIT on low-complexity datasets. The contig N50s of linked-read assemblies were higher than those of short-read assemblies but lower than those of long-read assemblies. Athena demonstrated a higher contig N50 than cloudSPAdes and generated the highest *#NC* among all of the assemblers for the metagenomic sequencing data from human stool samples. Long-read assemblers demonstrated the highest contig N50 but generated smaller *#MQ* and *#HQ* than short- and linked-read assemblers. metaFlye, Canu, and Lathe performed much better than the other long-read assemblers. metaFlye generated the highest *GF*s and *AL*s for both ONT and PacBio CLR datasets. Lathe produced a higher *#NC* than metaFlye and Canu using ONT data. Hybrid assemblies demonstrated comparable or lower contig N50s and generated higher *#HQ* and *#NC* than long-read assemblies. metaFlye-subassemblies (with “--subassemblies”) generated higher contig N50s and were more stable than the other hybrid assemblers.

**Figure 1.**
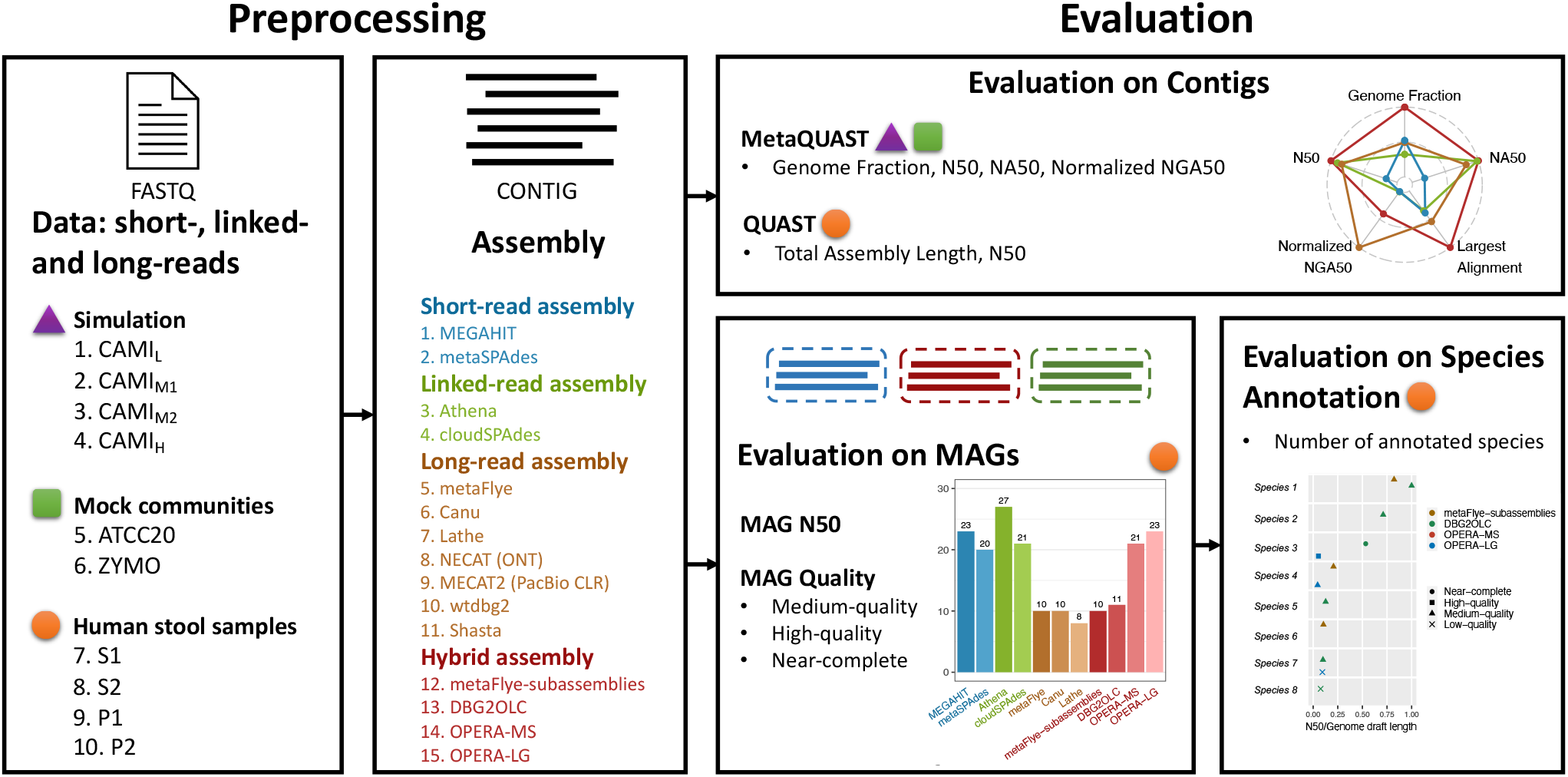
Data and workflow used to benchmark 15 metagenome assembly tools. **Preprocessing**: We collected 32 datasets generated by short-read, linked-read, and long-read sequencing from simulation, mock communities and four human stool samples, which were used to evaluate the performances of 15 de novo assembly tools. **Evaluation**: The following contig statistics were used to evaluate the assemblies: total assembly length (*AL*), genome fraction (*GF*), contig N50, NA50, and normalized NGA50. The assemblies generated from the real datasets were evaluated by metagenome-assembled genome (MAG) N50s; the numbers of medium-quality (*#MQ*), high-quality (*#HQ*), and near-complete (*#NC*) MAGs; and the numbers of annotated species.

## Results

### Metagenomic sequencing datasets from simulation, mock communities, and human stool samples

In this study, 32 datasets were collected and generated using three sequencing technologies (**Methods**; Table 1 and Fig. 1): short-read (Illumina HiSeq and BGISEQ-500), linked-read (10x Chromium), and long-read (PacBio CLR and ONT) sequencing datasets. These comprised the following datasets: (1) Simulation datasets: four CAMI communities with low (CAMI_L_), medium (CAMI_M1_ and CAMI_M2_), and high (CAMI_H_) complexities consisting of available short-reads, simulated 10x linked-reads, and simulated ONT long-reads. We merged CAMI datasets with high complexity from five time points into CAMI_H_ to avoid insufficient read depth. The 10x linked-reads and ONT long-reads were simulated for the four CAMI communities (**Methods**). (2) Mock datasets from two mock communities: ATCC-MSA-1003^40^ (ATCC20, sequenced by Illumina HiSeq 2500, 10x Chromium, and PacBio CLR) and ZymoBIOMICSTM Microbial Community Standard II with log distribution^41^ (ZYMO, sequenced by Illumina HiSeq 1500, ONT GridION, and ONT PromethION). (3) Real datasets: four stool samples from human gut microbiome, denoted S1 and S2 (sequenced by Illumina HiSeq 2500, BGISEQ-500, 10x Chromium, and PacBio CLR; Supplementary Figure 1), and P1 and P2^19,29^ (sequenced by Illumina HiSeq 4000, 10x Chromium, and ONT).

**Table 1.** The simulation, mock, and real datasets used to evaluate the performance of metagenome assembly tools. The PacBio CLR dataset of ATCC20 was downsampled to 50% to avoid out of memory issues.

| Datasets |  | Sequencing Platforms | Sources | Amount of sequencing (G) | Sequencing Depth (X) | Average Read Length (bp) | Community composition |
| --- | --- | --- | --- | --- | --- | --- | --- |
| Simulation | CAMI<sub>L</sub> | Illumina HiSeq | CAMI I | 15.0 | 95 | 150 | 40 genomes and 20 circular elements<sup>37</sup> |
|  |  | 10x Chromium | Simulation | 15.0 | 95 | 150 |  |
|  |  | ONT | Simulation | 15.0 | 95 | 4,194 |  |
|  | CAMI<sub>M1</sub> | Illumina HiSeq | CAMI I | 15.0 | 26 | 150 | 132 genomes and 100 circular elements<sup>37</sup> |
|  |  | 10x Chromium | Simulation | 15.0 | 26 | 150 |  |
|  |  | ONT | Simulation | 15.0 | 26 | 4,217 |  |
|  | CAMI<sub>M2</sub> | Illumina HiSeq | CAMI I | 15.0 | 26 | 150 | 132 genomes and 100 circular elements<sup>37</sup> |
|  |  | 10x Chromium | Simulation | 15.0 | 26 | 150 |  |
|  |  | ONT | Simulation | 15.0 | 26 | 4,186 |  |
|  | CAMI<sub>H</sub> | Illumina HiSeq | CAMI I | 75.0 | 27 | 150 | 596 genomes and 478 circular elements<sup>37</sup> |
|  |  | 10x Chromium | Simulation | 75.0 | 27 | 150 |  |
|  |  | ONT | Simulation | 75.0 | 27 | 4,112 |  |
| Mock | ATCC20 | Illumina HiSeq 2500 | SRR8359173 | 1.3 | 19 | 125 | 20 strains<sup>40</sup> |
|  |  | 10x Chromium | SRR12283286 | 108.7 | 1,622 | 150 |  |
|  |  | PacBio CLR | SRR12371719 | 253.5 | 3,784 | 8,394 |  |
|  | ZYMO | Illumina HiSeq 1500 | ERR2935805 | 9.7 | 133 | 101 | 10 species<sup>41</sup> |
|  |  | ONT GridION | ERR3152366 | 16.5 | 226 | 4,501 |  |
|  |  | ONT PromethION | ERR3152367 | 153.7 | 2,105 | 4,446 |  |
| Real | S1 | Illumina HiSeq 2500 | This study | 11.6 | 29 | 150 | Human gut microbiome |
|  |  | BGISeq-500 | This study | 16.8 | 42 | 100 |  |
|  |  | 10x Chromium | This study | 58.7 | 147 | 150 |  |
|  |  | PacBio CLR | This study | 6.3 | 16 | 8,878 |  |
|  | S2 | Illumina HiSeq 2500 | This study | 11.4 | 34 | 150 | Human gut microbiome |
|  |  | BGISeq-500 | This study | 15.0 | 44 | 100 |  |
|  |  | 10x Chromium | This study | 56.1 | 165 | 150 |  |
|  |  | PacBio CLR | This study | 8.4 | 25 | 8,973 |  |
|  | P1 | Illumina HiSeq 4000 | SRR6788327, SRR6807561 | 76.6 | 383 | 150 | Human gut microbiome |
|  |  | 10x Chromium | SRR6760786 | 35.8 | 179 | 150 |  |
|  |  | ONT MinION | SRR8427258 | 11.4 | 57 | 2,838 |  |
|  | P2 | Illumina HiSeq 4000 | SRR6788328, SRR6807555 | 77.6 | 776 | 150 | Human gut microbiome |
|  |  | 10x Chromium | SRR6760782 | 32.6 | 326 | 150 |  |
|  |  | ONT MinION | SRR8427257 | 6.1 | 61 | 1,800 |  |

### Metagenome assembly on short-read sequencing

We compared the assembly performance of metaSPAdes and MEGAHIT, the two mainstream short-read assemblers, on Illumina short-read sequencing data. The contigs generated by these two tools had comparable *GF*s (Fig. 2 a-f) and *AL*s (Fig. 3 a-d) on all datasets but nevertheless exhibited unique characteristics on different datasets. MEGAHIT showed better performance on the datasets with deeper sequencing depth (>100X; Table 1), such as ZYMO (133X), P1 (383X), and P2 (776X). For these datasets, the contigs of MEGAHIT had substantially higher N50s than those of metaSPAdes by 205.90% (ZYMO; Fig. 2 f), 148.60% (P1; Fig. 3 c), and 130.35% (P2; Fig. 3 d), respectively. By breaking the contigs at misassemblies, NA50 of MEGAHIT was 2.08 times higher than that of metaSPAdes on ZYMO (NA50: MEGAHIT = 167.51 kbp, metaSPAdes = 80.58 kbp; Fig. 2 f; Supplementary Table 1). By comparing the contig continuities of known species in ZYMO, we found that MEGAHIT also obtained 1.33 times higher normalized NGA50 than metaSPAdes (Fig. 2 f; Supplementary Figure 2). We grouped the contigs from the real datasets into MAGs and classified them based on different qualities (**Methods**). MEGAHIT produced higher *#MQ* and *#HQ* on P1 and P2 (*#MQ* in P1 and P2: MEGAHIT = 21, metaSPAdes = 6; *#HQ* in P1 and P2: MEGAHIT = 4, metaSPAdes = 1; Fig. 4 i-j, m-n) and achieved significantly higher MAG N50s than metaSPAdes (Wilcoxon rank-sum test p-value: P1 = 2.79e-5, P2 = 2.36e-3; Fig. 4 l and p; Supplementary Figure 3-4; Supplementary Table 2; **Methods**). We annotated the MAGs into species with a rigorous quality control process (**Methods**). In P1 and P2, the MAGs from MEGAHIT were annotated to seven species, whereas those from metaSPAdes were annotated to only two species (Fig. 5 c-d). metaSPAdes obtained much better contig continuity than MEGAHIT from the low- and medium-complexity datasets that were not deeply sequenced (<100X), such as CAMI_L_, CAMI_M1_, CAMI_M2_, and ATCC20. Its contig N50s, NA50s, and normalized NGA50s were, on average, 1.62, 1.66 and 1.43 times higher than those of MEGAHIT, respectively (Fig. 2 a-c and e; Supplementary Figure 5-6).

**Figure 2.**
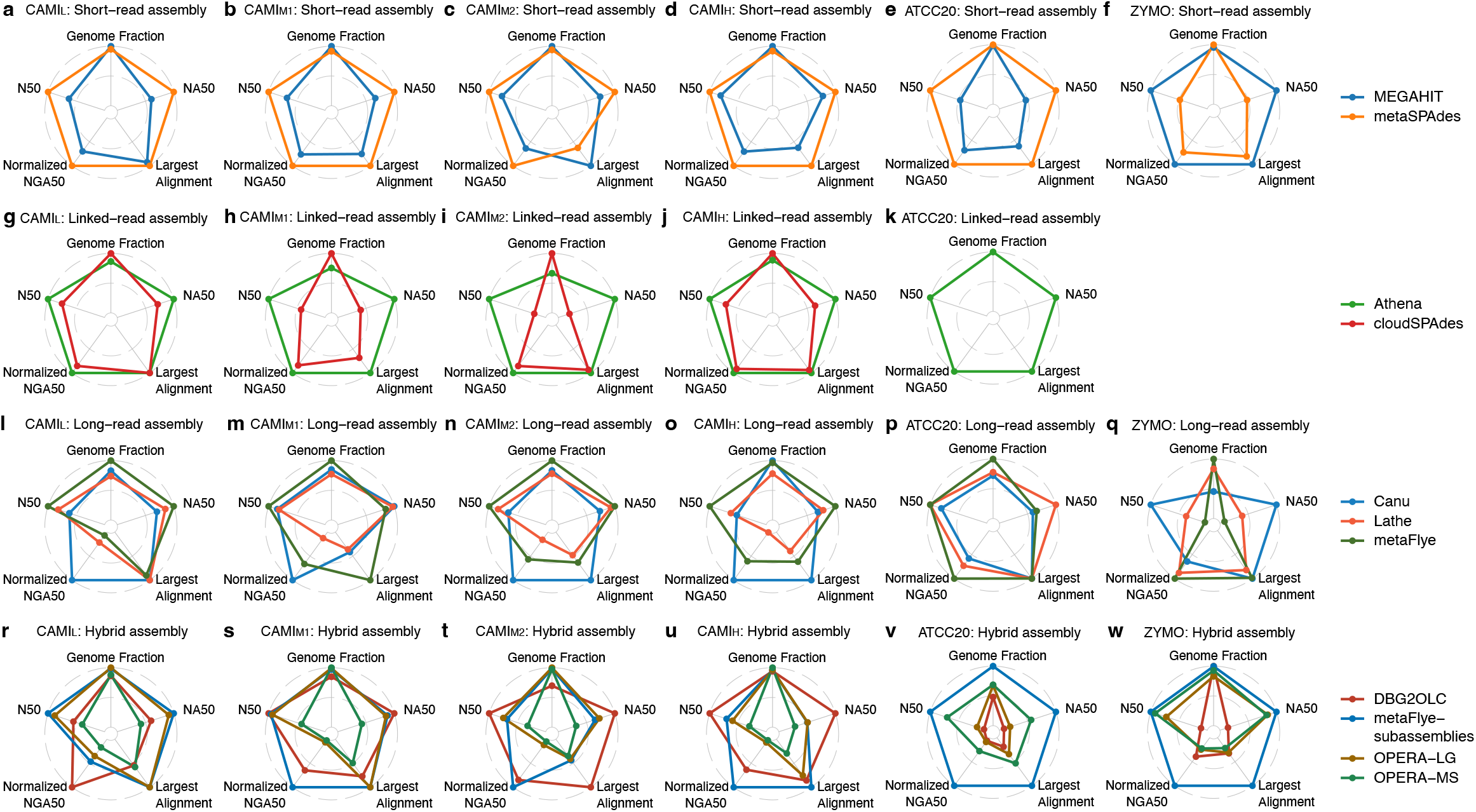
Contig statistics for the assemblies generated from the simulation and mock datasets (a-f for short-read assemblies; g-k for linked-read assemblies; l-q for long-read assemblies; r-w for hybrid assemblies). The long-reads used in q and w were sequenced by ONT GridION.

**Figure 3.**
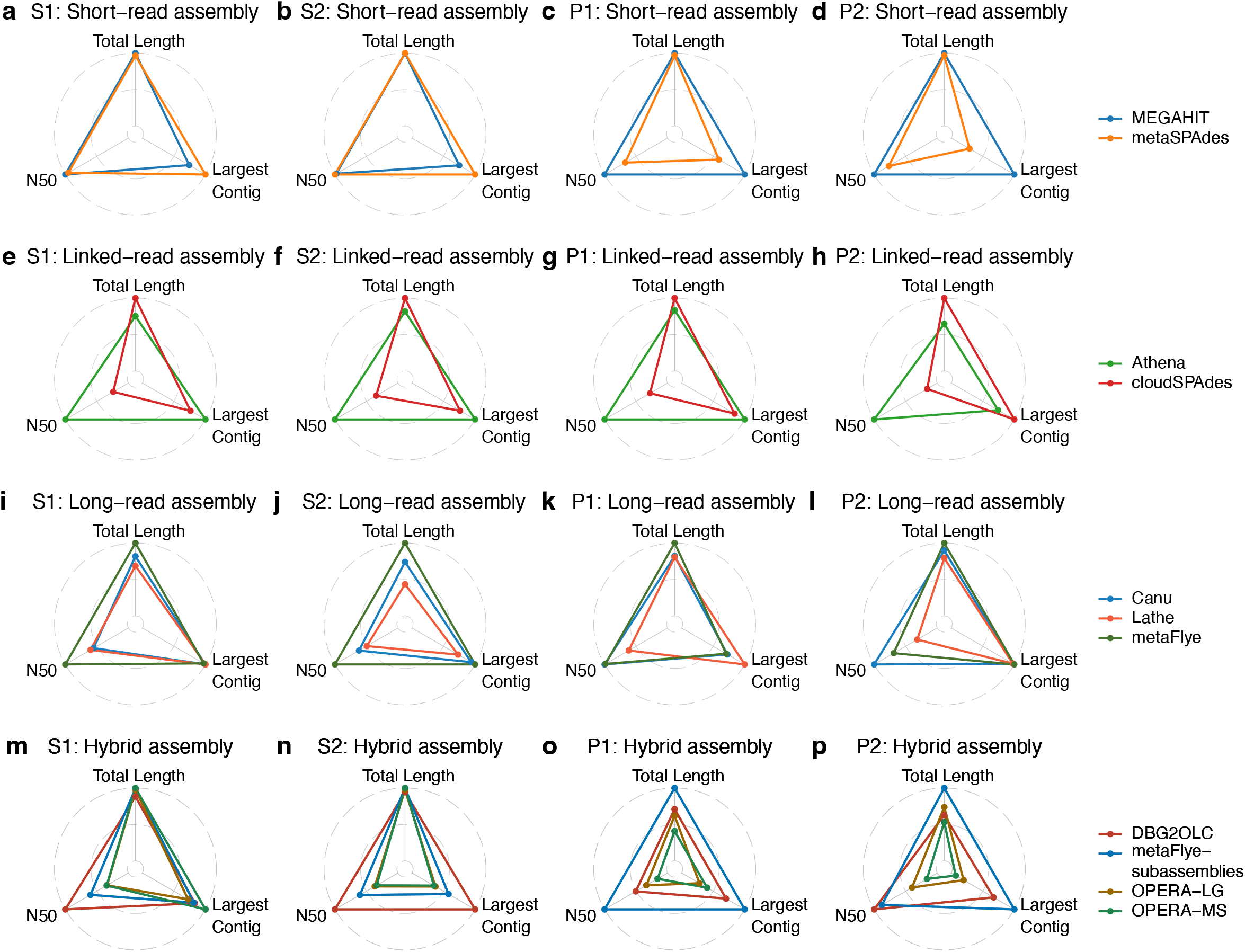
Contig statistics for the assemblies generated from real datasets (a-d for short-read assemblies; e-h for linked-read assemblies; i-l for long-read assemblies; m-p for hybrid assemblies). The short-reads used in a and b were generated by Illumina HiSeq 2500.

**Figure 4.**
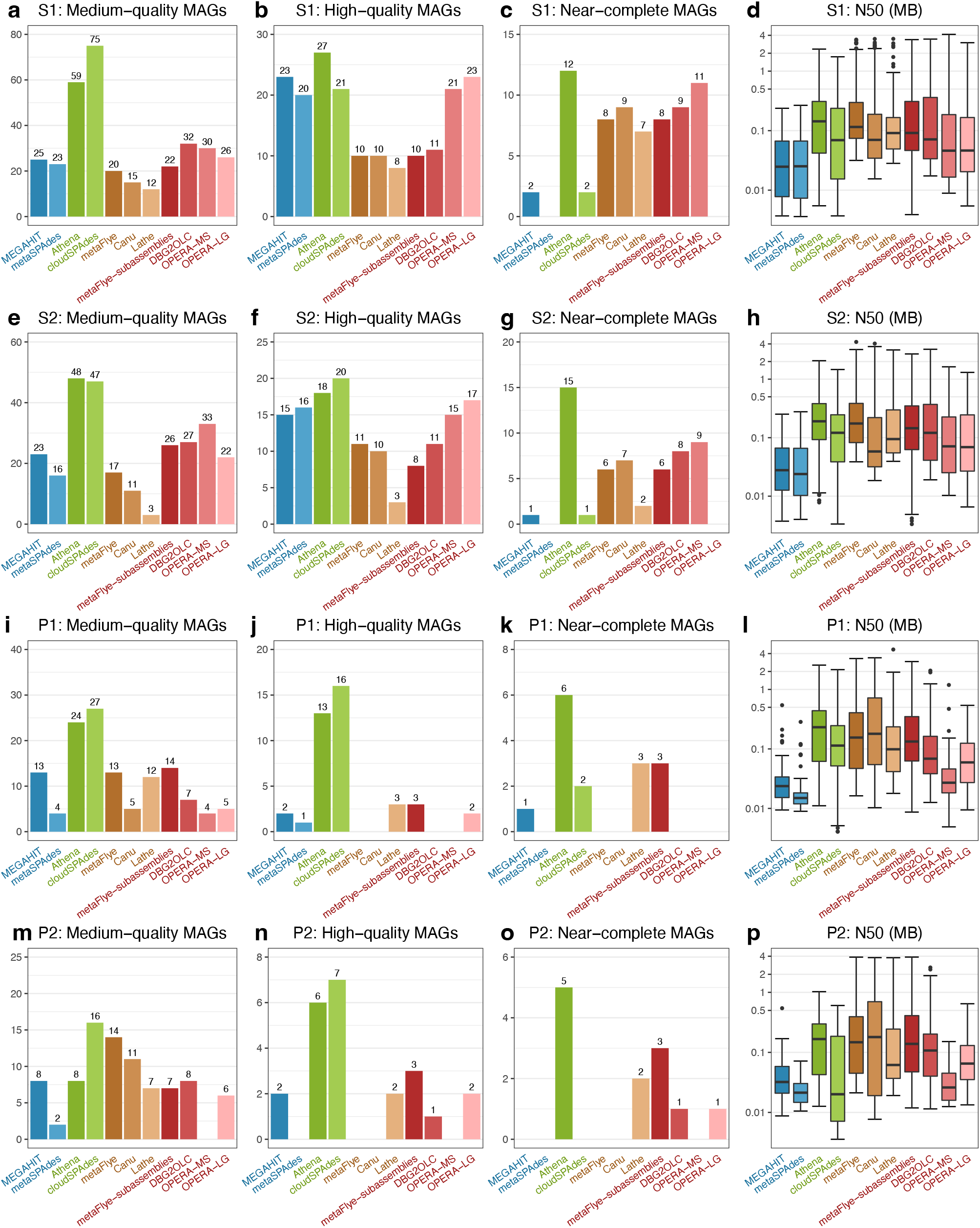
Numbers of medium-quality, high-quality, and near-complete metagenome-assembled genomes (MAGs) and the MAG N50 values for the assemblies generated from the real datasets (a-d for S1; e-h for S2; i-l for P1; m-p for P2). The short-reads used in a-d and e-h were generated by Illumina HiSeq 2500.

By comparing the assemblies using the datasets from Illumina HiSeq 2500 and BGISEQ-500, we observed MEGAHIT and metaSPAdes showed a similar trend that the two platforms generated comparable contig contiguity, *#MQ, #HQ* (Supplementary Table 1), and numbers of annotated species on S1 and S2 (Supplementary Figure 7). Their MAGs could be annotated to unique species. For example on S1 (Supplementary Figure 7), *Sutterella faecalis* and *Ruminococcus bicirculans* were uniquely annotated from the metaSPAdes assemblies on Illumina HiSeq 2500 and BGISEQ-500, respectively.

### Metagenome assembly on linked-read sequencing

We compared the assemblies generated by cloudSPAdes and Athena on 10x linked-read sequencing data, which revealed that cloudSPAdes generated higher *GF*s than Athena on the simulation datasets (127.69% on average; Fig. 2 g-j) and produced higher *AL*s on the real datasets (average *AL*: cloudSPAdes = 343.65 Mbp, Athena = 267.55 Mbp; Fig. 3 e-h; Supplementary Table 1). cloudSPAdes failed to assemble the ATCC20 linked-read dataset due to insufficient memory (>1TB RAM); similar trends of *GF*s on the ATCC20 dataset were also reported in a previous study^18^. In the assessments using simulation datasets, the contigs from Athena had higher N50s (250.55% on average), NA50s (261.32% on average), and normalized NGA50s (115.53% on average) than those from cloudSPAdes (Fig. 2 g-j; Supplementary Figure 5). Tolstoganov et al. also reported higher contig N50s and NA50s from Athena on ATCC20^18^. A similar trend was observed for the real datasets: contig N50s (average N50: Athena = 158.92 kbp, cloudSPAdes = 43.16 kbp; Fig. 3 e-h; Supplementary Table 1) and MAG N50s (Wilcoxon rank-sum test p-value: S1 = 2.04e-5, S2 = 3.05e-6, P1 = 4.86e-4, P2 = 5.01e-5; Fig. 4 d, h, l and p; Supplementary Figure 3-4 and 8-9; Supplementary Table 2) from Athena were significantly higher than those from cloudSPAdes. Athena also generated substantially more *#NC* than cloudSPAdes from the real datasets (*#NC* in total: Athena = 38, cloudSPAdes = 5; Fig. 4 c, g, k, and o). A comparable number of species was annotated from the MAGs generated by the two assembly tools, and both tools identified unique species (Fig. 5 e-h). For example, Athena identified three unique species (*Roseburia intestinalis, Phascolarctobacterium faecium*, and *Desulfovibrio fairfieldensis*), while cloudSPAdes identified four unique species (*Sutterella faecalis, Bifidobacterium adolescentis, Flavonifractor plautii*, and a *Longibaculum sp*.) on the S1 dataset (Fig. 5 e).

**Figure 5.**
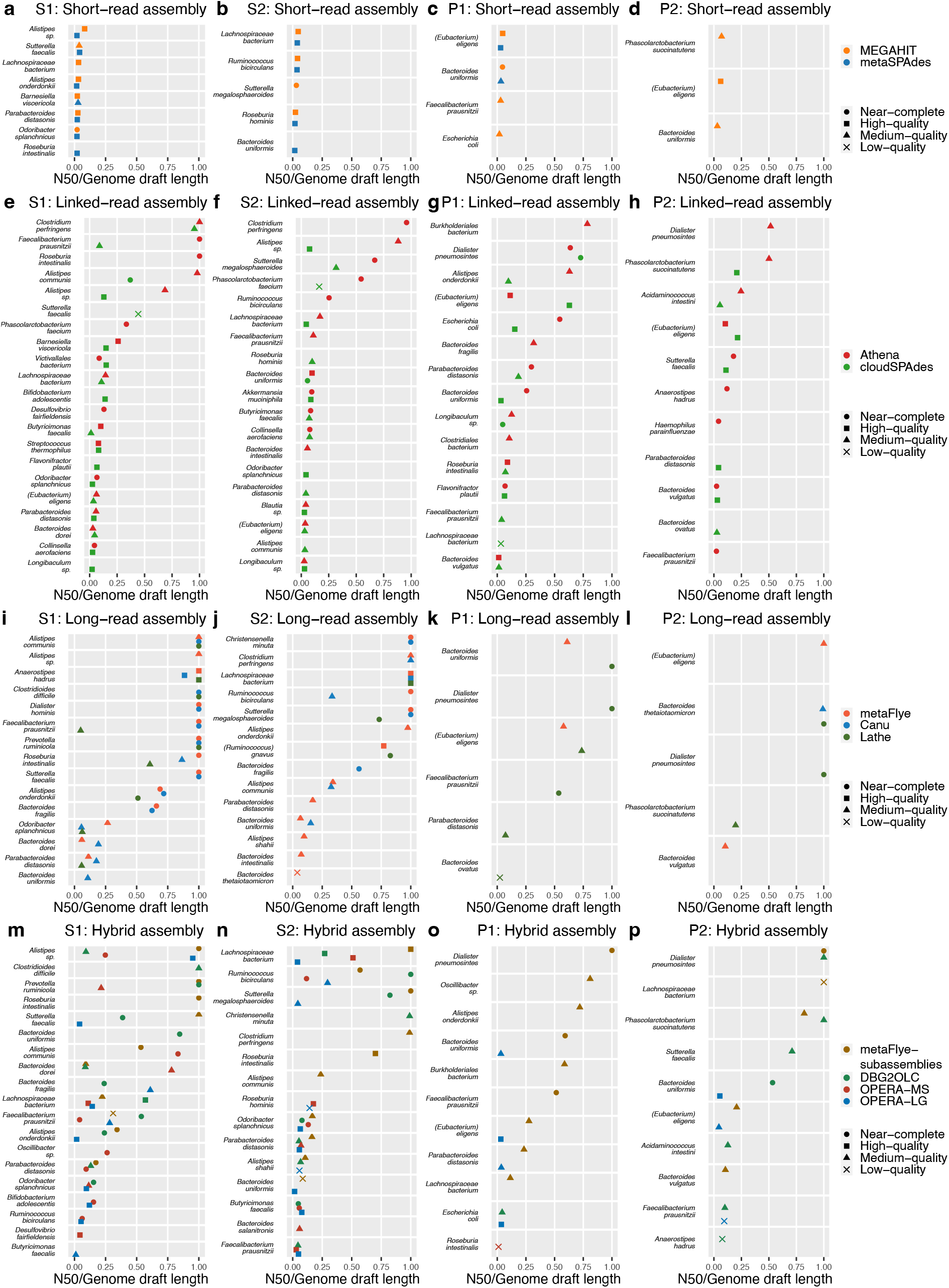
MAG annotations of the assemblies generated from the real datasets (a-d for short-read assemblies; e-h for linked-read assemblies; i-l for long-read assemblies; m-p for hybrid assemblies). The N50/genome draft length was used to evaluate the continuity of MAGs. The short-reads used in a and b were sequenced by Illumina HiSeq 2500.

### Metagenome assembly on long-read sequencing

We compared the performances of seven long-read assembly tools using PacBio CLR and ONT data: Shasta, wtdbg2, MECAT2 (PacBio CLR only), NECAT (ONT only), Canu, metaFlye, and Lathe. Canu, metaFlye, and Lathe generated assemblies with significantly higher *GF*s on the simulation and mock datasets (328.45 times on average; Supplementary Table 1) and the assemblies with much longer *AL*s on all of the real datasets (4.64 times on average; except wtdbg2; Supplementary Table 1) than the other four tools (Shasta, wtdbg2, MECAT2, and NECAT). This may be because Shasta, wtdbg2, MECAT2, and NECAT were not designed for metagenome assembly and may preferentially generate long contigs derived from a single species. Therefore, we only included Canu, metaFlye, and Lathe in the subsequent analysis as they generated assemblies with reasonable lengths.

The contigs generated by metaFlye had higher *GF*s than those generated by Canu and Lathe on most of the simulation and mock datasets (1.44 times on average, except Canu on CAMI_H_; Fig. 2 l-q). metaFlye also generated the highest *AL*s on the real datasets (an average of 1.40 times greater than Canu and Lathe; Fig. 3 i-l). The contig continuities of metaFlye assemblies were better than those of the corresponding Lathe assemblies on most datasets sequenced by either PacBio CLR or ONT: metaFlye generated higher contig N50s (1.60 times), NA50s (1.27 times), and normalized NGA50s (79.39 times) than Lathe on CAMI_H_ (Fig. 2 o; Supplementary Figure 5), and higher contig N50s (average contig N50 on S1, S2 and P1: metaFlye = 199.04 kbp, Lathe = 112.63 kbp; Fig. 3 i-k; Supplementary Table 1) and significantly higher MAG N50s (Wilcoxon rank-sum test p-value: S1 = 2.27e-4, S2 = 9.85e-4, P1 = 4.83e-2; Fig. 4 d, h and l; Supplementary Figure 3, 8-9; Supplementary Table 2) than Lathe on the real datasets (S1, S2, and P1).

Compared to the assemblies generated by metaFlye and Lathe, the assemblies generated by Canu had higher (or comparable) contig continuity on ONT data, but lower (or comparable) continuity on PacBio CLR data. Eight datasets were generated by ONT, including CAMI_L_, CAMI_M1_, CAMI_M2_, CAMI_H_, ZYMO (GridION and PromethION), P1, and P2. Canu generated assemblies with higher normalized NGA50s (4.77 times on average) than metaFlye on the four CAMI datasets and generated assemblies with higher contig N50s (16.40 times on average) and NA50s (8.89 times on average) than metaFlye on the GridION and PromethION datasets from ZYMO (Fig. 2 l-o and q; Supplementary Figure 5; Supplementary Table 1). Canu also generated assemblies with higher contig continuity than those generated by metaFlye and Lathe on P1 and P2 (average contig N50s on P1 and P2: Canu = 233.62 kbp, metaFlye = 189.79 kbp, Lathe = 102.91 kbp; Fig. 3 k-l; Supplementary Table 1). Regarding the datasets sequenced by PacBio CLR (ATCC20, S1, and S2), metaFlye and Lathe produced assemblies with substantially higher contig N50s (124.33% on average), NA50s (139.95% on average), and normalized NGA50s (149.04% on average) than Canu on ATCC20 (Fig. 2 p; Supplementary Figure 10); metaFlye produced assemblies with higher contig N50s (171.32% on average) and MAG N50s (Wilcoxon rank-sum test p-value: S1 = 7.45e-5, S2 = 3.77e-8) than Canu on S1 and S2 (Fig. 3 i-j; Fig. 4 d and h; Supplementary Figure 8-9; Supplementary Table 2).

We further evaluated the MAG qualities and annotations of the assemblies generated by these three tools. For the PacBio CLR datasets from S1 and S2, metaFlye and Canu generated comparable *#HQ* and *#NC*, which were much larger than those generated by Lathe (*#HQ*: metaFlye = 21, Canu = 20, Lathe = 11; *#NC*: metaFlye = 14, Canu = 16, Lathe = 9; Fig. 4 b-c, f-g). They also generated more MAGs that were annotated to species (metaFlye = 26, Canu = 22, Lathe = 12; Fig. 5 i-j). For the ONT datasets from P1 and P2, the opposite trend was observed in the MAG qualities (*#HQ*: Lathe = 5, Canu = 0, metaFlye = 0; *#NC*: lathe = 5, metaFlye = 0, Canu = 0; Fig. 4 j-k, n-o) and annotated species (Lathe = 9, metaFlye = 4, Canu = 1; Fig. 5 k-l). As the core assembly algorithms of Lathe are the same as those of Canu and metaFlye, the above observations imply that parameter optimization and sequencing error-correction are vital for ONT data assembly. We also found that all the three assembly tools identified uniquely annotated species, e.g., metaFlye, Canu, and Lathe uniquely reported *Alistipes sp*. (S1), *Bacteroides uniformis* (S1), and *Dialister pneumosintes* (P1), respectively (Fig. 5 i and k).

### Metagenome hybrid assembly on short- and long-read sequencing

We evaluated the performance of four hybrid assembly tools, namely, metaFlye-subassemblies (**Methods**), DBG2OLC, OPERA-LG, and OPERA-MS, on short- and long-read sequencing datasets. Compared with the other tools, metaFlye-subassemblies produced assemblies with significantly higher *GF* (167.16% on average; Fig. 2 v) on ATCC20 and significantly higher *AL* on P1 (182.44% on average; Fig. 3 o) and P2 (160.87% on average; Fig. 3 p). The four hybrid assembly tools generated comparable *GF*s and *AL*s on all of the other datasets (Fig. 2 r-u and w; Fig. 3 m-n). metaFlye-subassemblies and DBG2OLC generated assemblies with substantially higher contig continuity than OPERA-MS and OPERA-LG on most datasets. For example, metaFlye-subassemblies produced assemblies with higher contig N50s (2.26 times on average), NA50s (2.61 times on average), and normalized NGA50s (4.92 times on average; Supplementary Figure 11-13) than OPERA-LG and OPERA-MS on ATCC20 and ZYMO (Fig. 2 v-w). metaFlye-subassemblies also obtained assemblies with higher contig N50s (2.97 times on average; Fig. 3 m-p) and MAG N50s than OPERA-LG and OPERA-MS (the Wilcoxon rank-sum test p-values were S1: OPERA-LG = 3.27e-5 and OPERA-MS = 1.65e-4; S2: OPERA-LG = 1.60e-4 and OPERA-MS = 1.05e-5; P1: OPERA-LG = 8.02e-7 and OPERA-MS = 9.90e-14; P2: OPERA-LG = 2.37e-3 and OPERA-MS = 1.79e-10; Fig. 4 d, h, l and p; Supplementary Figure 3-4, 8-9; Supplementary Table 2) on all of the real datasets. Furthermore, metaFlye-subassemblies had higher contig N50s (27.99 times on average), NA50s (19.18 times on average), and normalized NGA50s (6.84 times on average) than DBG2OLC assemblies on the low-complexity mock datasets such as ATCC20 and ZYMO (Fig. 2 v-w; Supplementary Figure 11-13; Supplementary Table 1).

We observed inconsistent trends in the MAG qualities of hybrid assemblies from different sequencing platform combinations. OPERA-MS and OPERA-LG generated assemblies with higher *#HQ* than those generated by the other tools on S1 and S2 (*#HQ*: OPERA-MS = 36, OPERA-LG = 40, metaFlye-subassemblies = 18, DBG2OLC = 22; Fig. 4 b and f), which were sequenced by Illumina and PacBio CLR. OPERA-MS assemblies showed higher *#NC* values than those generated by OPERA-LG (OPERA-MS = 20, OPERA-LG=0; Fig. 4 c and g) from these two samples. For P1 and P2 sequenced by Illumina and ONT, metaFlye-subassemblies obtained assemblies with higher *#HQ* and *#NC* than the other tools (*#HQ*: metaFlye-subassemblies = 6, DBG2OLC = 1, OPERA-MS = 0, OPERA-LG = 4; *#NC*: metaFlye-subassemblies = 6, DBG2OLC = 1, OPERA-MS = 0, OPERA-LG = 1; Fig. 4 j-k, n-o). The four assembly tools identified comparable numbers of annotated species on S1 and S2 (metaFlye-subassemblies = 20, DBG2OLC = 21, OPERA-MS = 20, OPERA-LG = 20; Fig. 5 m-n). metaFlye-subassemblies identified more annotated species than the other three tools on P1 and P2 (metaFlye-subassemblies = 14, DBG2OLC = 8, OPERA-MS = 1, OPERA-LG = 7; Fig. 5 o-p). The MAGs generated by various tools were also annotated to unique species, e.g., *R. intestinalis, Clostridioides difficile, Desulfovibrio fairfiendensis*, and *Butyricimonas faecalis* were uniquely identified from the assemblies of metaFlye-subassemblies, DBG2OLC, OPERA-MS, and OPERA-LG on S1 (Fig. 5 m).

### Metagenome assembly on different sequencing strategies

We compared the best assembly statistics (e.g. *AL, GF, N50, #HQ*, and the number of annotated species; Fig. 6-7; Supplementary Table 3) of the four sequencing strategies discussed above: (1) short-read sequencing, (2) linked-read sequencing, (3) long-read sequencing, and (4) hybrid sequencing (short- and long-read sequencing). We observed the *GF*s and *AL*s of hybrid assemblies were higher than or comparable to those generated from short-read or long-read assemblies for all datasets (Fig. 6). The contig continuities of hybrid and linked-read assemblies were lower than or comparable with long-read assemblies, and significantly better than those from short-read assemblies (Fig. 6).

**Figure 6.**
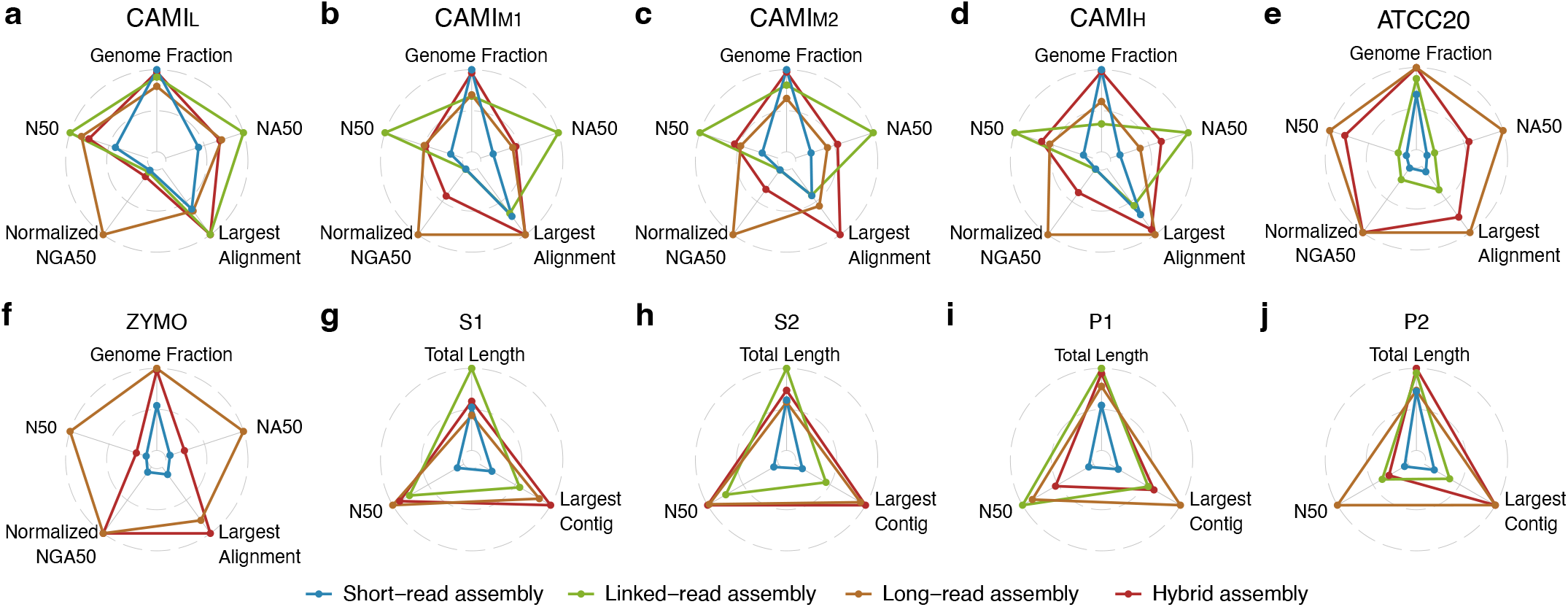
Contig statistics for the assemblies generated from simulation datasets (a-d); mock datasets (e-f); and real datasets (g-j).

**Figure 7.**
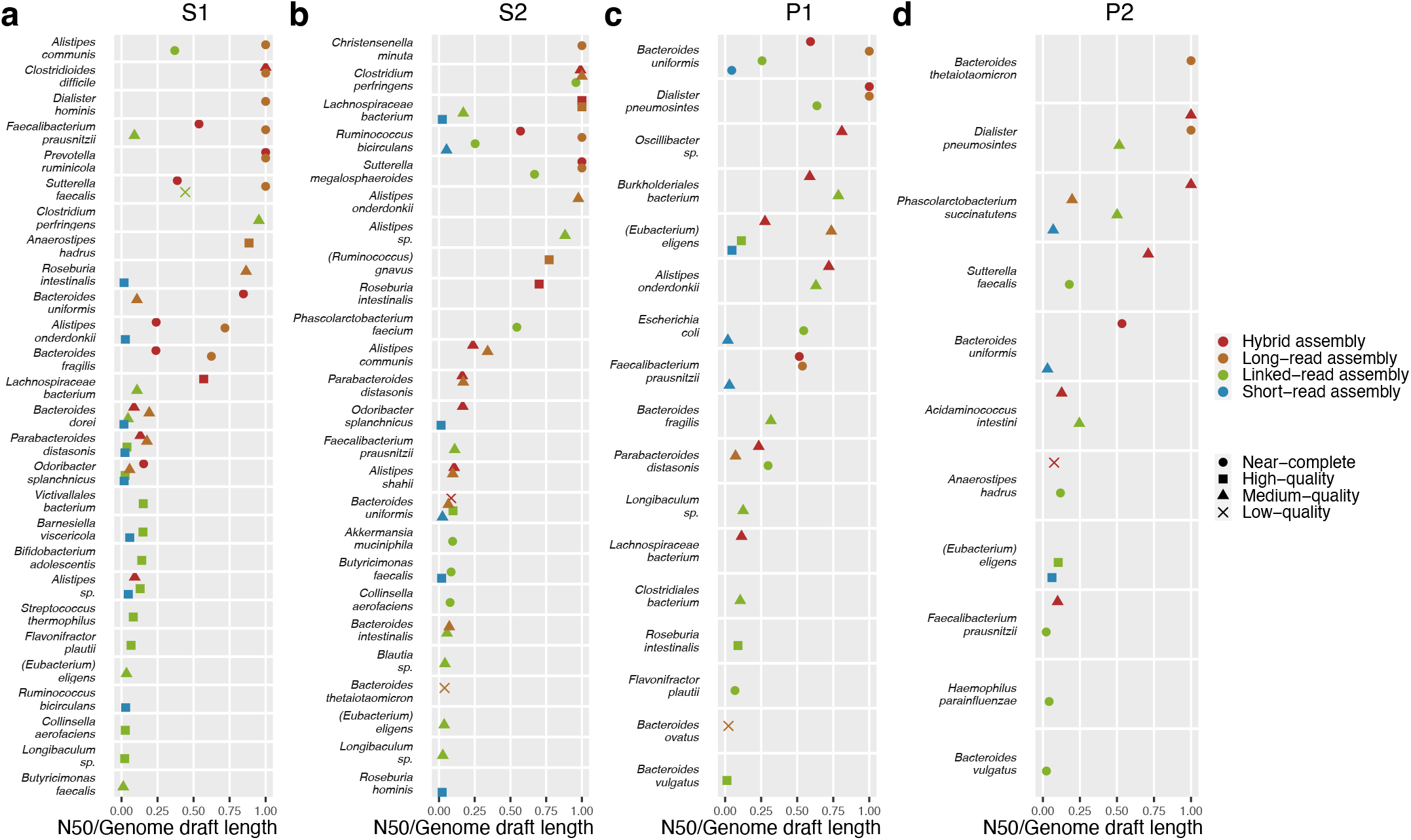
MAG annotations of the assemblies generated from real datasets with different sequencing strategies.

Nevertheless, linked-read assemblies had the highest *#MQ* (166), *#HQ* (70), *#NC* (38) and identified the highest number of annotated species (55) in all of the real datasets (Fig. 7; Supplementary Table 3). These values were much higher than those of the short-read assemblies (*#MQ* = 77, *#HQ* = 43, *#NC* = 4, number of annotated species = 21; Fig. 7; Supplementary Table 3), the long-read assemblies (*#MQ* = 64, *#HQ* = 26, *#NC* = 21, number of annotated species = 36; Fig. 7; Supplementary Table 3), and the hybrid assemblies (*#MQ* = 87, *#HQ* = 46, *#NC* = 26, number of annotated species = 38; Fig.7; Supplementary Table 3). Notably, the assembly generated from each type of sequencing strategy could identify unique species, implying that these assemblies were somewhat complementary; for example, *R. hominis, P. faecium, Alistipes onderdonkii*, and *R. intestinalis* were uniquely annotated by the short-read, linked-read, long-read, and hybrid assemblies on the S2 dataset, respectively (Fig. 7 b).

### Evaluation of computational time and resources

We compared the running time (real time) and peak memory consumption (maximum resident set size [RSS]) of the assembly tools applied to the simulation datasets (Fig. 8). We found that metaSPAdes had a longer running time (1.92 times on average) and substantially larger memory usage (8.72 times on average) than MEGAHIT for short-read assembly. cloudSPAdes had a significantly longer running time (5.80 times on average) and consumed higher peak memory (8.73 times on average) than Athena for linked-read assembly. Canu required more than 7 days to complete the metagenome assembly on any of the CAMI datasets, which was more than twice as long as the other long-read assembly tools (we excluded Canu in Fig. 8 because it exceeded our server wall-clock time). Of the other long-read assemblers, Lathe and metaFlye had the highest peak memory consumption (3.57 times on average) and the longest running time (5.82 times on average), respectively (except for CAMI_H_). Of the hybrid assembly tools, DBG2OLC (6.80 times slower than the other tools on average) and metaFlye-subassemblies (9.30 times faster than the other tools on average) were the slowest and fastest tools, respectively. OPERA-MS and OPERA-LG had substantially higher peak-memory consumption than metaFlye-subassemblies and DBG2OLC (2.58 times on average). In addition, the trends in the CPU times recorded for all of these assembly tools were the same as the trends in the real times (Supplementary Figure 14).

**Figure 8.**
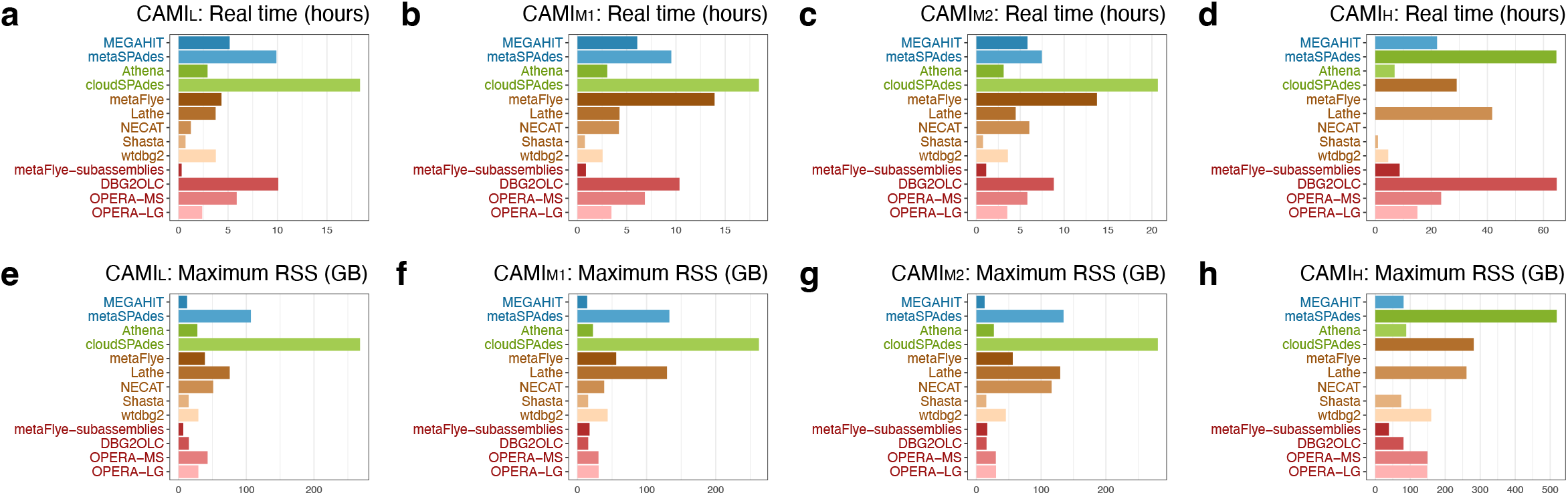
Computational resources (real time and resident set size [RSS]) consumed by the assembly tools in analyzing CAMI datasets. metaFlye and NECAT were not used for the analysis of CAMI_H_ because it was found that they exceeded the maximum memory limitation.

## Discussion

Metagenome assembly is the most straightforward way to identify the “microbial dark matter” from the microbiota and is therefore a core analytical method. Many metagenome assembly tools have been developed to analyze the data from various sequencing technologies, but there lacks an independent, comprehensive, and up-to-date investigation of these tools. We benchmarked the performance of the 15 most widely used assemblers on simulation, mock, and real datasets with diverse complexities and provided practical guidance for tool selection.

Recent studies^42–44^ have demonstrated the applicability of representing microbial genomes using MAGs generated from large short-read metagenomic sequencing cohorts. Nevertheless, the short-read assembly usually generates highly fragmented contigs and poor quality MAGs. By comparing MEGAHIT and metaSPAdes, we observed that MEGAHIT outperformed metaSPAdes in generating assemblies from deeply sequenced datasets (>100x), probably because MEGAHIT has an optimized data structure and algorithms to analyze large datasets. However, metaSPAdes performed better than MEGAHIT in generating assemblies from low-complexity datasets, which is consistent with the observations from a previous study^38^.

Linked-reads are short-reads that are labeled with barcodes to facilitate the linking of short-reads if they are derived from the same long DNA fragments. We found that linked-read assemblies had consistently better contig continuity than short-read assemblies but sometimes worse continuity than long-read assemblies. This is probably because linked-reads help to resolve the ambiguous branches and circles from repetitive sequences in assembly graphs^18,19^. However, linked-reads nevertheless fail to capture the tandem repeats and highly variable regions presented within long fragments. Furthermore, we observed that Athena generated assemblies containing contigs with higher continuity and higher *#NC* than those generated by cloudSPAdes, although cloudSPAdes generated more sequences. The *#NC*s from linked-read assemblies were higher than those from long-read assemblies, which may be attributable to the high read depth and the low base-error rate of linked-read sequencing.

Long-read assemblers can generate the most continuous contigs, but individual contigs usually do not represent circular microbial genomes and sometimes cannot be grouped into high-quality MAGs. This may be because (1) high-molecular-weight DNA cannot be easily extracted from some microbes^19^; (2) high sequencing costs prevent sufficient long-reads being obtained; and (3) error-prone long-reads result in low-quality MAGs. In this study, we compared the performances of seven state-of-the-art long-read assemblers and found that Canu, metaFlye, and Lathe performed substantially better than the other assemblers. metaFlye generated assemblies with the highest *GF* and *AL*, consistent with the observation of a previous study^39^. Lathe generated assemblies with significantly more *#NC* than metaFlye and Canu from the ONT datasets, suggesting that sequencing error-correction is an essential step to improve MAG quality generated from ONT long-reads.

Hybrid assembly has been proposed as a way to correct assembly errors from error-prone long-reads by adding short-reads with high base quality. In this study, the hybrid assemblies had comparable or lower contig continuity than those from long-read assemblers, but the former had higher *#NC*. This observation is in line with our expectations, as the short-reads were mainly used to reduce misassemblies rather than fill gaps between contigs. We also observed that the contig continuities from overlap-layout-consensus tools (metaFlye-subassemblies and DBG2OLC) were better than those generated by *de Bruijn* graph tools (OPERA-LG and OPERA-MS).

Reconstructing the genomes of low- (0.1%-1%) and ultra-low (<0.1%) abundance species is a challenging task^45,46^. We evaluated the performances of different assemblers in assembling the genomes of species with low- and ultra-low abundance in ATCC20 (Supplementary Table 4). Short-read assemblers were unable to assemble any genomes of low-abundance species (GF>50%; Supplementary Figure 6), whereas all of these species were identified by linked-read (Athena; Supplementary Table 4), long-read (metaFlye and Shasta; Supplementary Figure 10), and hybrid assemblers (metaFlye-subassemblies and DBG2OLC; Supplementary Figure 11). For the five low-abundance species in ATCC20, the long-read (metaFlye) and hybrid (metaFlye-subassemblies) assemblers generated higher contig continuity than Athena on the linked-reads (Supplementary Table 4). Only one (ATCC_8482) of the five ultra-low abundance species was obtained by long-read (Lathe and metaFlye; Supplementary Figure 10) and hybrid (metaFlye-subassembly; Supplementary Figure 11) assemblers, suggesting that ultra-low abundance species are still difficult to be assembled even if long-reads have been adopted.

Recently, PacBio HiFi technology has shown its great success in deciphering microbial communities^47^. We also compared the long-read assemblers on the PacBio HiFi dataset from ATCC20. Similar to our findings for PacBio CLR, we observed that metaFlye, Lathe, and Canu generated substantially higher *GF*s than Shasta, wtdbg2 and MECAT2 (8.98 times on average, Supplementary Table 1). metaFlye and Lathe obtained significantly higher N50s (2.50 times on average), NA50s (2.06 times on average), and normalized NGA50s (1.37 times on average) than Canu (Supplementary Table 1, Supplementary Figure 15). We found that the long-read assemblers on PacBio HiFi and CLR sequencing platforms produced comparable *GF*s, N50s, NA50s, and normalized NGA50s (Supplementary Table 1). The misassemblies generated by Canu on PacBio CLR reads were substantially higher (2.17 times on average) than on PacBio HiFi reads (Supplementary Table 1).

In this study, we thoroughly investigated the pros and cons of the metagenome assembly tools using datasets generated by a range of sequencing technologies – short-read, long-read, linked-read, and hybrid sequencing – and we provide practical guidelines to assist end-users to select the best strategy for their purposes. We believe that our findings will be invaluable to the microbiome research community and will shed light on future genome-based microbiome-wide association studies.

## Methods

### Simulate linked-reads and ONT long-reads for CAMI datasets

We simulated 10x linked-reads and ONT long-reads using LRTK-SIM (v201912229)^48^ and CAMISIM (v1.2-beta)^49^ given the taxonomic composition in CAMI datasets. The simulated total nucleotides were the same as those of the available four short-read CAMI datasets.

### Sample preparation and sequencing for S1 and S2

The S1 and S2 datasets were obtained from two subjects with a typical Chinese diet and who had not taken any antibiotics, probiotics, or prebiotics in the three months prior to the sample collection. Their stool samples were collected, aliquoted, and stored at -80°C until analysis. Total microbial DNA was extracted using the QIAamp DNA stool mini kit (Qiagen, Valencia, CA, USA) according to the manufacturer’s protocol. For short-read sequencing, fecal microbial DNA of the two subjects was sequenced by paired-end sequencing with a coverage of more than 30 million reads per sample using Illumina HiSeq 2500 (Illumina, CA, USA) and BGISEQ-500 (BGI, ShenZhen, China), respectively. For linked-read sequencing, we followed the strategies described by Bishara et al.^19^ for library preparation on a 10x Chromium System (10x Genomics, CA, USA) and performed sequencing on an Illumina HiSeq 2500 (Illumina, CA, USA). For long-read sequencing, the SMRTbell libraries were prepared with the 20-kb Template Preparation using BluePippinTM Size selection system (15-kb size cutoff) protocol and were then sequenced in SMRT cells (Pacific Biosciences, CA, USA) with magnetic bead loading and P4-C2 or P6-C4 chemistry.

### Metagenome assembly

We used MEGAHIT (v1.2.9)^14^ and metaSPAdes (v3.15.0)^11^ for short-read assembly; cloudSPAdes (v3.12.0-dev)^18^ and Athena (v1.3)^19^ for linked-read assembly; Shasta (v0.7.0)^32^, wtdbg2 (v2.5)^33^, MECAT2 (v20190314)^30^, NECAT (v0.01)^31^, metaFlye (v2.8.3)^24^, Canu (v2.1.1)^26^, and Lathe (v20210210)^29^ for long-read assembly; and DBG2OLC (v20200724)^34^, and metaFlye (v2.8.3)^24^, OPERA-LG (v2.0.6)^35^, and OPERA-MS (v0.8.3)^36^ for hybrid assembly. To enable the hybrid mode of metaFlye (metaFlye-subassemblies), we combined the contigs assembled from short-reads (metaSPAdes) and long-reads (metaFlye) using the “--subassemblies” option. We used “--pacbio-hifi” for metaFlye HiFi assembly^47^.

Most of the assembly tools were run using default parameters, but we adjusted the parameters in the following cases to avoid out of memory issues: (1) metaFlye on CAMI_H_ was run with “flye --asm-coverage 50”; and (2) NECAT on CAMI_H_ was run with “MIN_READ_LENGTH=8000”. All commands are available at https://github.com/ZhangZhenmiao/metagenome_assembly. The running times (user, system, and real times) and the maximum peak memory (RSS) consumptions of the assembly tools were retrieved using the Linux command “/usr/bin/time -v”. All of the assembly tools were run on Linux machines with a dual 64-core AMD EPYC 7742 2.25GHz base clock speed 256MB L3 cache CPU with 1 TB memory.

### Contig statistics

We generated *AL*, contig N50, and MAG N50 by QUAST (v5.0.2)^50^ after removing the contigs shorter than 1 kb. We enabled the MetaQUAST mode^51^ to obtain contig NA50 and NGA50 for each species from the datasets for which reference genomes were available. We used the parameter “--fragmented --min-alignment 500 --unique-mapping” in MetaQUAST to disable ambiguous alignments. To compare NGA50s across different samples, we defined normalized NGA50 by averaging NGA50/genome size across all of the species in the sample.

### Contig binning and MAG qualities

To prepare the inputs of MetaBat2^52^ for contig binning, we used BWA (v0.7.17)^53^ and minimap2 (v2.17)^54^ to align short-reads (or linked-reads) and long-reads to the contigs, respectively. Minimap2 used the parameters “-ax map-pb” and “-ax map-ont” to align PacBio CLR and ONT reads, respectively. For hybrid assembly, the short-read alignment was adopted as the input of MetaBat2. The alignment file was sorted by coordinates using SAMtools (v1.9)^55^, and the contig coverage was extracted by the “jgi_summarize_bam_contig_depths” program in MetaBat2. MetaBat2 (v2.12.1)^52^ was used to group the contigs into MAGs using both contig coverage and sequence characteristics. We explored the single-copy gene completeness and contamination of each MAG using CheckM (v1.1.2)^56^. The transfer RNAs (tRNAs) and ribosomal RNAs (5S, 16S, and 23S rRNAs) were detected by ARAGORN (v1.2.38)^57^ and barrnap (v0.9)^58^, respectively. MAGs were defined as high-quality (completeness > 90%, contamination < 5%), medium-quality (completeness ≥ 50%, contamination < 10%), or low-quality (otherwise). Near-complete MAGs were those high-quality MAGs with 5S, 16S, and 23S rRNAs, and at least 18 tRNAs^59^.

### Annotate MAGs into species

We removed poorly assembled MAGs with contig N50s < 50 kbp, completeness < 75%, or contamination > 25%, and annotated the contigs in MAGs with Kraken2 (v2.1.2)^60^. To determine the dominant species identified for each MAG, we used “assign_species.py” from “metagenomics_workflows”^61^, which has been adopted in previous studies^19,29^. dRep^62^ was used to remove the redundant MAGs from the same species.

## Data availability

The CAMI short-reads were downloaded from “1st CAMI Challenge Dataset 1 CAMI_low”, “1st CAMI Challenge Dataset 2 CAMI_medium” and “1st CAMI Challenge Dataset 3 CAMI_high” of CAMI challenge website. The other available datasets were downloaded from NCBI Sequence Read Archive (SRA). The Illumina HiSeq 2500 short-reads, 10x linked-reads and PacBio CLR long-reads of ATCC20 were available with the accession codes SRR8359173, SRR12283286 and SRR12371719, respectively. We also used the PacBio HiFi long-reads of ATCC20 from SRR9202034 and SRR9328980. The Illumina HiSeq 1500 short-reads, ONT GridION and ONT PromethION long-reads of ZYMO were collected from ERR2935805, ERR3152366 and ERR3152367, respectively. The Illumina HiSeq 4000 short-reads (P1: SRR6788327, SRR6807561; P2: SRR6788328, SRR6807555), 10x linked-reads (P1: SRR6760786 ; P2: SRR6760782) and ONT long-reads (P1: SRR8427258; P2: SRR8427257) of P1 and P2 were also downloaded from SRA. The Illumina HiSeq 2500 short-reads, 10x linked-reads, and long-reads of S1 and S2 are available in SRA (PRJNA841170).

## Acknowledgements

The authors thank the Research Grants Council of Hong Kong, Hong Kong Baptist University, and HKBU Research Committee for their kind support of this project.

## Author contributions statement

LZ conceived the study; ZMZ conducted the experiments and analyzed the results; ZMZ and LZ wrote the manuscript; CY reviewed the manuscript; and XDF helped with the library preparation and sequencing of S1 and S2. All of the authors have read and approved the final manuscript.

## Supplementary information

**Supplementary Notes**: Supplementary Figure 1-15.

**Supplementary Figure 1**: Species composition and Shannon diversity of the S1 and S2 datasets obtained from different sequencing platforms.

**Supplementary Figure 2**: Contig continuities and genome fractions (*GF*) for the short-read assemblies on the ZYMO dataset. The red and green colors on the y-axis represent low- and ultra-low abundance species, respectively. The suffixes in the figure legend indicate the corresponding sequencing platforms.

**Supplementary Figure 3**: MAG N50s and coverage distribution for P1. The suffixes in the figure legend indicate the corresponding sequencing platforms. The green, blue, red, and black colors denote short-read, linked-read, long-read, and hybrid assembly tools, respectively.

**Supplementary Figure 4**: MAG N50s and coverage distribution for P2. The suffixes in the figure legend indicate the corresponding sequencing platforms. The green, blue, red, and black colors denote short-read, linked-read, long-read, and hybrid assembly tools, respectively.

**Supplementary Figure 5**: Distribution of normalized NGA50s for the species in the four CAMI datasets. The suffixes in the figure legend indicate the corresponding sequencing platforms. The green, blue, red, and black colors denote short-read, linked-read, long-read, and hybrid assembly tools, respectively.

**Supplementary Figure 6**: Contig continuities and genome fractions (*GF*) for the short-read assemblies on the ATCC20 dataset. The red and green colors on the y-axis represent low- and ultra-low abundance species, respectively. The suffixes in the figure legend indicate the corresponding sequencing platforms.

**Supplementary Figure 7**: The MAG annotations for S1 and S2 generated by BGISEQ-500 and Illumina HiSeq 2500.

**Supplementary Figure 8**: MAG N50s and coverage distribution for S1. The suffixes in the figure legend indicate the corresponding sequencing platforms. The green, blue, red, and black colors denote short-read, linked-read, long-read, and hybrid assembly tools, respectively.

**Supplementary Figure 9**: MAG N50s and coverage distribution for S2. The suffixes in the figure legend indicate the corresponding sequencing platforms. The green, blue, red, and black colors denote short-read, linked-read, long-read, and hybrid assembly tools, respectively.

**Supplementary Figure 10**: Contig continuities and genome fractions (*GF*) for the long-read assemblies on the PacBio CLR dataset of ATCC20. The red and green colors on the y-axis represent low- and ultra-low abundance species, respectively. The suffixes in the figure legend indicate the corresponding sequencing platforms.

**Supplementary Figure 11**: Contig continuities and genome fractions (*GF*) for the hybrid assemblies on the ATCC20 dataset. The red and green colors on the y-axis represent low- and ultra-low abundance species, respectively. The suffixes in the figure legend indicate the corresponding sequencing platforms.

**Supplementary Figure 12**: Contig continuities and genome fractions (*GF*) for the hybrid assemblies (Illumina + ONT GridION) on the ZYMO dataset. The red and green colors on the y-axis represent low- and ultra-low abundance species, respectively. The suffixes in the figure legend indicate the corresponding sequencing platforms.

**Supplementary Figure 13**: Contig continuities and genome fractions (*GF*) for the hybrid assemblies (Illumina + ONT PromethION) on the ZYMO dataset. The red and green colors on the y-axis represent low- and ultra-low abundance species, respectively. The suffixes in the figure legend indicate the corresponding sequencing platforms.

**Supplementary Figure 14**: CPU time consumed by the assembly tools in processing the CAMI datasets. We removed metaFlye and NECAT on CAMI_H_ because they exceeded the maximum memory limitation.

**Supplementary Figure 15**: Contig continuities and genome fractions (*GF*) for the long-read assemblies on the PacBio HiFi dataset of ATCC20. The red and green colors on the y-axis represent low- and ultra-low abundance species, respectively. The suffixes in the figure legend indicate the corresponding sequencing platforms.

**Supplementary Table 1**: QUAST report of the assembly results generated from all of the datasets, and the *#MQ, #HQ*, and *#NC* for the real datasets.

**Supplementary Table 2**: The p-values (Wilcoxon rank-sum tests) of MAG N50s generated by different assembly tools on the real datasets.

**Supplementary Table 3**: The best assembly statistics of the assemblies from different sequencing technologies.

**Supplementary Table 4**: NGA50s for the low- and ultra-low abundance species from the ATCC20 dataset.

## References

1. Korem, T. et al. Growth dynamics of gut microbiota in health and disease inferred from single metagenomic samples. Science 349, 1101–1106 (2015).

2. Xu, P. & Gunsolley, J. Application of metagenomics in understanding oral health and disease. Virulence 5, 424–432 (2014).

3. Martín, R., Miquel, S., Langella, P. & Bermúdez-Humarán, L. G. The role of metagenomics in understanding the human microbiome in health and disease. Virulence 5, 413–423 (2014).

4. Simon, C. & Daniel, R. Metagenomic analyses: past and future trends. Appl. environmental microbiology 77, 1153–1161 (2011).

5. Browne, H. P. et al. Culturing of ‘unculturable’human microbiota reveals novel taxa and extensive sporulation. Nature 533, 543–546 (2016).

6. Forster, S. C. et al. A human gut bacterial genome and culture collection for improved metagenomic analyses. Nat. biotechnology 37, 186–192 (2019).

7. Singh, B. K. Exploring microbial diversity for biotechnology: the way forward. Trends biotechnology 28, 111–116 (2010).

8. Yang, C. et al. A review of computational tools for generating metagenome-assembled genomes from metagenomic sequencing data. Comput. Struct. Biotechnol. J. 19, 6301–6314 (2021).

9. Peng, Y., Leung, H. C., Yiu, S.-M. & Chin, F. Y. Meta-idba: a de novo assembler for metagenomic data. Bioinformatics 27, i94–i101 (2011).

10. Peng, Y., Leung, H. C., Yiu, S.-M. & Chin, F. Y. Idba-ud: a de novo assembler for single-cell and metagenomic sequencing data with highly uneven depth. Bioinformatics 28, 1420–1428 (2012).

11. Nurk, S., Meleshko, D., Korobeynikov, A. & Pevzner, P. A. metaspades: a new versatile metagenomic assembler. Genome research 27, 824–834 (2017).

12. Bankevich, A. et al. Spades: a new genome assembly algorithm and its applications to single-cell sequencing. J. computational biology 19, 455–477 (2012).

13. Prjibelski, A. D. et al. Exspander: a universal repeat resolver for dna fragment assembly. Bioinformatics 30, i293–i301 (2014).

14. Li, D., Liu, C.-M., Luo, R., Sadakane, K. & Lam, T.-W. Megahit: an ultra-fast single-node solution for large and complex metagenomics assembly via succinct de bruijn graph. Bioinformatics 31, 1674–1676 (2015).

15. Besser, J., Carleton, H. A., Gerner-Smidt, P., Lindsey, R. L. & Trees, E. Next-generation sequencing technologies and their application to the study and control of bacterial infections. Clin. microbiology infection 24, 335–341 (2018).

16. Zlitni, S. et al. Strain-resolved microbiome sequencing reveals mobile elements that drive bacterial competition on a clinical timescale. Genome medicine 12, 1–17 (2020).

17. Roodgar, M. et al. Longitudinal linked-read sequencing reveals ecological and evolutionary responses of a human gut microbiome during antibiotic treatment. Genome research 31, 1433–1446 (2021).

18. Tolstoganov, I., Bankevich, A., Chen, Z. & Pevzner, P. A. cloudspades: assembly of synthetic long reads using de bruijn graphs. Bioinformatics 35, i61–i70 (2019).

19. Bishara, A. et al. High-quality genome sequences of uncultured microbes by assembly of read clouds. Nat. biotechnology 36, 1067–1075 (2018).

20. Chin, C.-S. et al. Nonhybrid, finished microbial genome assemblies from long-read smrt sequencing data. Nat. methods 10, 563–569 (2013).

21. Loman, N. J., Quick, J. & Simpson, J. T. A complete bacterial genome assembled de novo using only nanopore sequencing data. Nat. methods 12, 733–735 (2015).

22. Tsai, Y.-C. et al. Resolving the complexity of human skin metagenomes using single-molecule sequencing. MBio 7, e01948–15 (2016).

23. Koren, S. et al. Reducing assembly complexity of microbial genomes with single-molecule sequencing. Genome biology 14, 1–16 (2013).

24. Kolmogorov, M. et al. metaflye: scalable long-read metagenome assembly using repeat graphs. Nat. Methods 17, 1103–1110 (2020).

25. Kolmogorov, M., Yuan, J., Lin, Y. & Pevzner, P. A. Assembly of long, error-prone reads using repeat graphs. Nat. biotechnology 37, 540–546 (2019).

26. Koren, S. et al. Canu: scalable and accurate long-read assembly via adaptive k-mer weighting and repeat separation. Genome research 27, 722–736 (2017).

27. Myers, E. W. et al. A whole-genome assembly of drosophila. Science 287, 2196–2204 (2000).

28. Miller, J. R. et al. Aggressive assembly of pyrosequencing reads with mates. Bioinformatics 24, 2818–2824 (2008).

29. Moss, E. L., Maghini, D. G. & Bhatt, A. S. Complete, closed bacterial genomes from microbiomes using nanopore sequencing. Nat. biotechnology 38, 701–707 (2020).

30. Xiao, C.-L. et al. Mecat: fast mapping, error correction, and de novo assembly for single-molecule sequencing reads. nature methods 14, 1072–1074 (2017).

31. Chen, Y. et al. Efficient assembly of nanopore reads via highly accurate and intact error correction. Nat. Commun. 12, 1–10 (2021).

32. Shafin, K. et al. Nanopore sequencing and the shasta toolkit enable efficient de novo assembly of eleven human genomes. Nat. biotechnology 38, 1044–1053 (2020).

33. Ruan, J. & Li, H. Fast and accurate long-read assembly with wtdbg2. Nat. methods 17, 155–158 (2020).

34. Ye, C., Hill, C. M., Wu, S., Ruan, J. & Ma, Z. S. Dbg2olc: efficient assembly of large genomes using long erroneous reads of the third generation sequencing technologies. Sci. reports 6, 1–9 (2016).

35. Gao, S., Bertrand, D., Chia, B. K. & Nagarajan, N. Opera-lg: efficient and exact scaffolding of large, repeat-rich eukaryotic genomes with performance guarantees. Genome biology 17, 1–16 (2016).

36. Bertrand, D. et al. Hybrid metagenomic assembly enables high-resolution analysis of resistance determinants and mobile elements in human microbiomes. Nat. biotechnology 37, 937–944 (2019).

37. Sczyrba, A. et al. Critical assessment of metagenome interpretation—a benchmark of metagenomics software. Nat. methods 14, 1063–1071 (2017).

38. Meyer, F. et al. Tutorial: assessing metagenomics software with the cami benchmarking toolkit. Nat. protocols 16, 1785–1801 (2021).

39. Latorre-Pérez, A., Villalba-Bermell, P., Pascual, J. & Vilanova, C. Assembly methods for nanopore-based metagenomic sequencing: a comparative study. Sci. reports 10, 1–14 (2020).

40. ATCC-MSA-1003. https://www.atcc.org/products/msa-1003.

41. Nicholls, S. M., Quick, J. C., Tang, S. & Loman, N. J. Ultra-deep, long-read nanopore sequencing of mock microbial community standards. Gigascience 8, giz043 (2019).

42. Parks, D. H. et al. Recovery of nearly 8,000 metagenome-assembled genomes substantially expands the tree of life. Nat. microbiology 2, 1533–1542 (2017).

43. Almeida, A. et al. A new genomic blueprint of the human gut microbiota. Nature 568, 499–504 (2019).

44. Almeida, A. et al. A unified catalog of 204,938 reference genomes from the human gut microbiome. Nat. biotechnology 39, 105–114 (2021).

45. Cleary, B. et al. Detection of low-abundance bacterial strains in metagenomic datasets by eigengenome partitioning. Nat. biotechnology 33, 1053–1060 (2015).

46. Luo, Y., Yu, Y. W., Zeng, J., Berger, B. & Peng, J. Metagenomic binning through low-density hashing. Bioinformatics 35, 219–226 (2019).

47. Bickhart, D. M. et al. Generating lineage-resolved, complete metagenome-assembled genomes from complex microbial communities. Nat. biotechnology 1–9 (2022).

48. Zhang, L., Zhou, X., Weng, Z. & Sidow, A. Assessment of human diploid genome assembly with 10x linked-reads data. Gigascience 8, giz141 (2019).

49. Fritz, A. et al. Camisim: simulating metagenomes and microbial communities. Microbiome 7, 1–12 (2019).

50. Gurevich, A., Saveliev, V., Vyahhi, N. & Tesler, G. Quast: quality assessment tool for genome assemblies. Bioinformatics 29, 1072–1075 (2013).

51. Mikheenko, A., Saveliev, V. & Gurevich, A. Metaquast: evaluation of metagenome assemblies. Bioinformatics 32, 1088–1090 (2016).

52. Kang, D. D. et al. Metabat 2: an adaptive binning algorithm for robust and efficient genome reconstruction from metagenome assemblies. PeerJ 7, e7359 (2019).

53. Li, H. Aligning sequence reads, clone sequences and assembly contigs with bwa-mem. arXiv preprint 1303.3997 (2013).

54. Li, H. Minimap2: pairwise alignment for nucleotide sequences. Bioinformatics 34, 3094–3100 (2018).

55. Li, H. et al. The sequence alignment/map format and samtools. Bioinformatics 25, 2078–2079 (2009).

56. Parks, D. H., Imelfort, M., Skennerton, C. T., Hugenholtz, P. & Tyson, G. W. Checkm: assessing the quality of microbial genomes recovered from isolates, single cells, and metagenomes. Genome research 25, 1043–1055 (2015).

57. Laslett, D. & Canback, B. Aragorn, a program to detect trna genes and tmrna genes in nucleotide sequences. Nucleic acids research 32, 11–16 (2004).

58. Seemann, T. barrnap. https://github.com/tseemann/barrnap (2018).

59. Bowers, R. M. et al. Minimum information about a single amplified genome (misag) and a metagenome-assembled genome (mimag) of bacteria and archaea. Nat. biotechnology 35, 725–731 (2017).

60. Wood, D. E., Lu, J. & Langmead, B. Improved metagenomic analysis with kraken 2. Genome biology 20, 1–13 (2019).

61. Moss, E. L. metagenomics_workflows. https://github.com/elimoss/metagenomics_workflows (2019).

62. Olm, M. R., Brown, C. T., Brooks, B. & Banfield, J. F. drep: a tool for fast and accurate genomic comparisons that enables improved genome recovery from metagenomes through de-replication. The ISME journal 11, 2864–2868 (2017).

