## Supplementary Notes for "Benchmarking *de novo* assembly methods on metagenomic sequencing data"

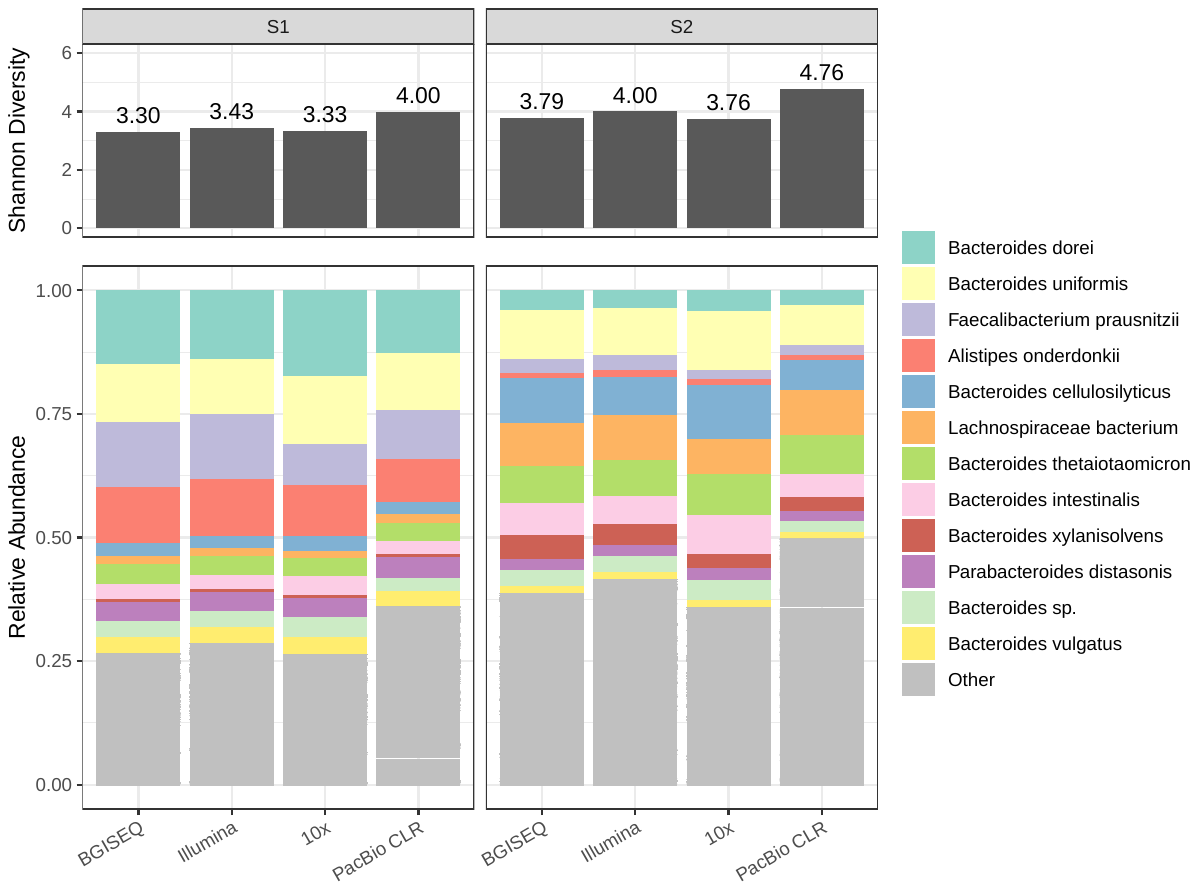

**Supplementary Figure 1.** Species composition and Shannon diversity of the S1 and S2 datasets obtained from different sequencing platforms.

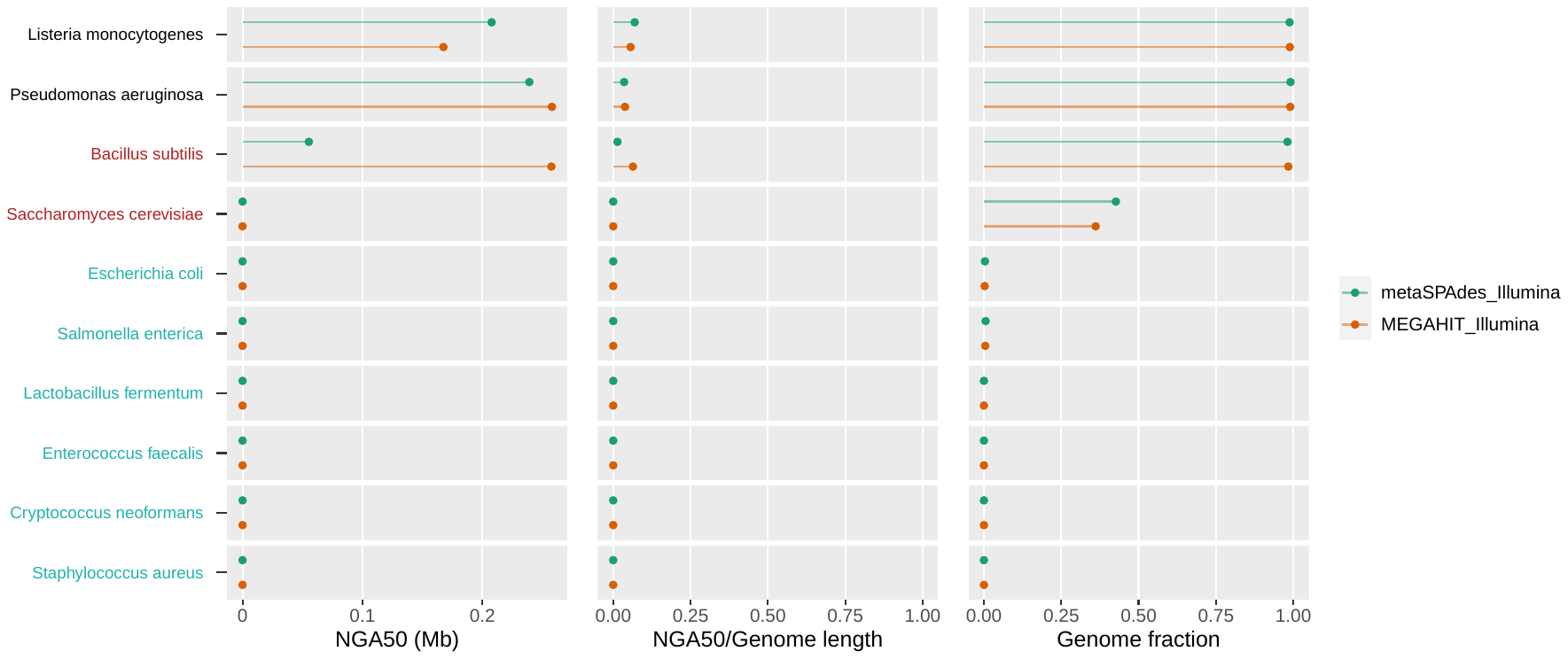

**Supplementary Figure 2.** Contig continuities and genome fractions (*GF*) for the short-read assemblies on the ZYMO dataset. The red and green colors on the y-axis represent low- and ultra-low abundance species, respectively. The suffixes in the figure legend indicate the corresponding sequencing platforms.

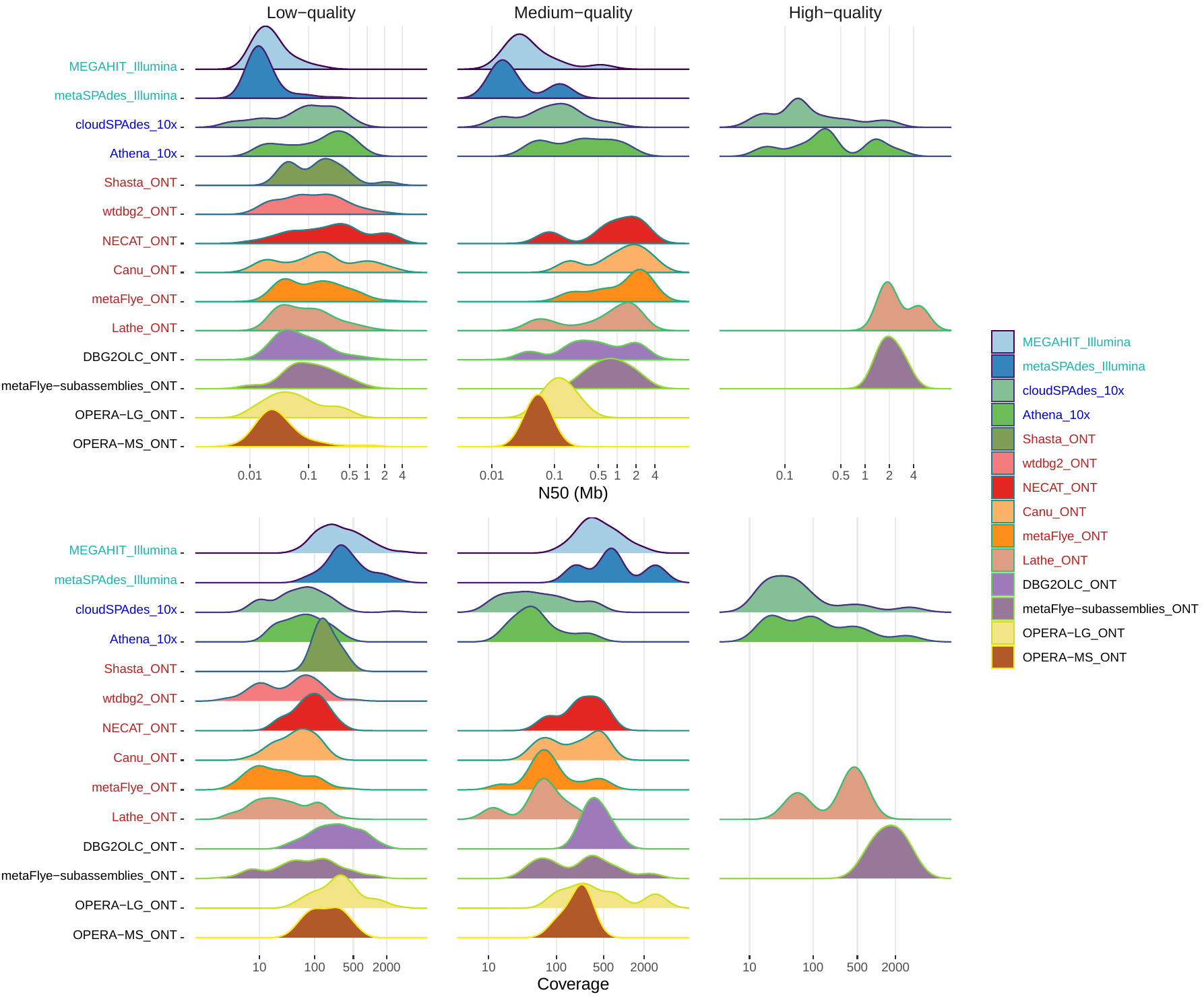

**Supplementary Figure 3.** MAG N50s and coverage distribution for P1. The suffixes in the figure legend indicate the corresponding sequencing platforms. The green, blue, red, and black colors denote short-read, linked-read, long-read, and hybrid assembly tools, respectively.

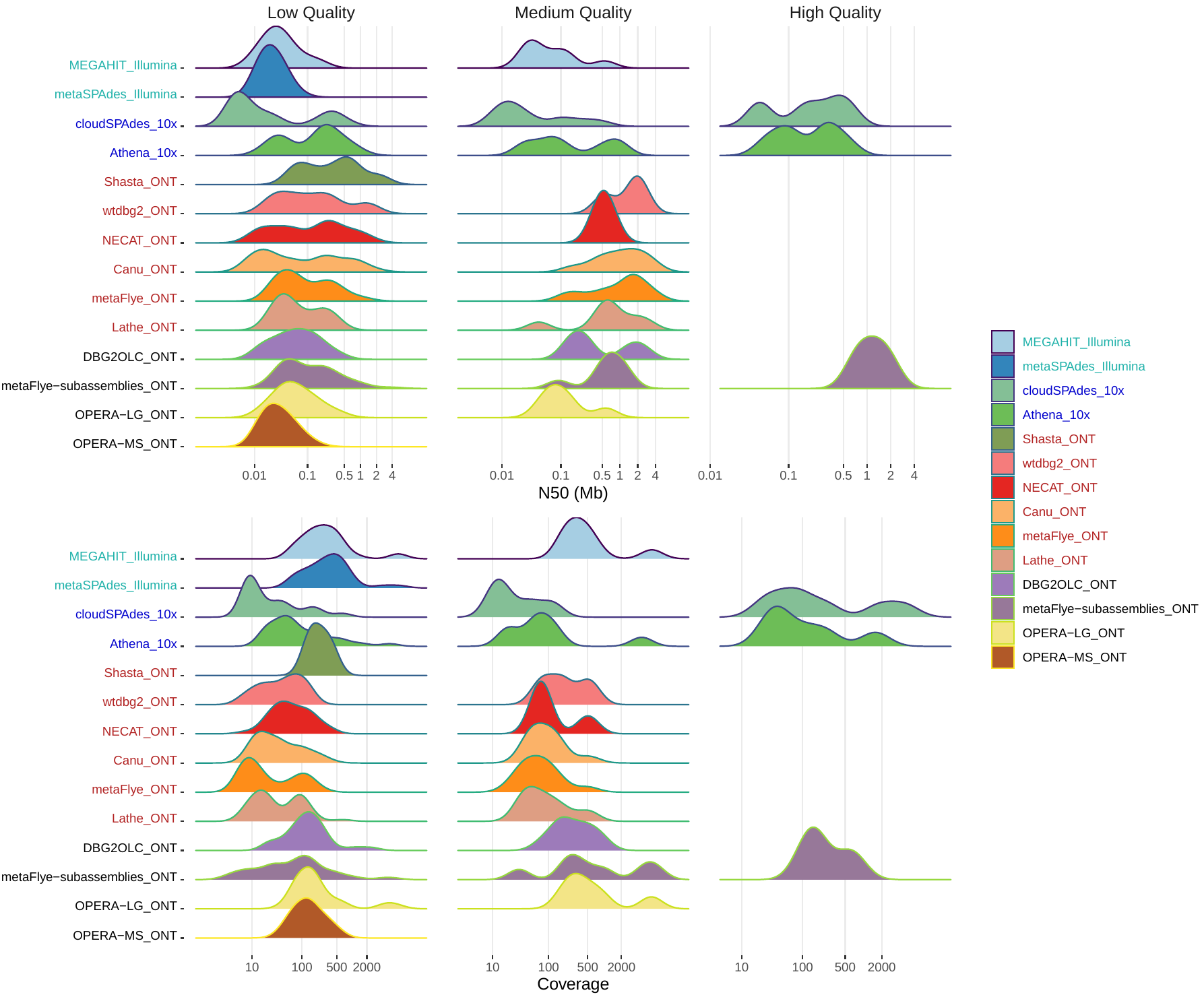

**Supplementary Figure 4.** MAG N50s and coverage distribution for P2. The suffixes in the figure legend indicate the corresponding sequencing platforms. The green, blue, red, and black colors denote short-read, linked-read, long-read, and hybrid assembly tools, respectively.

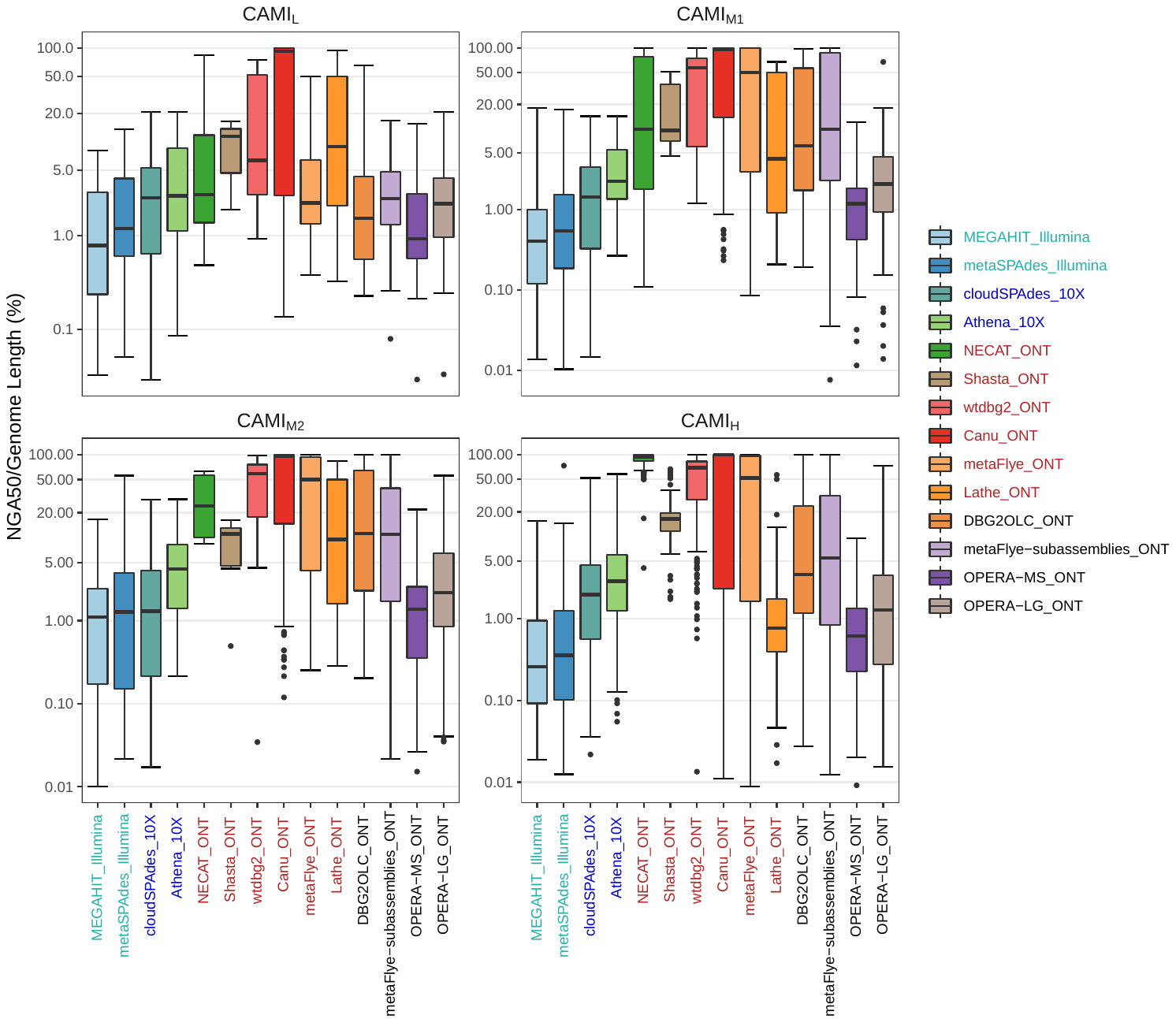

**Supplementary Figure 5.** Distribution of normalized NGA50s for the species in the four CAMI datasets. The suffixes in the figure legend indicate the corresponding sequencing platforms. The green, blue, red, and black colors denote short-read, linked-read, long-read, and hybrid assembly tools, respectively.

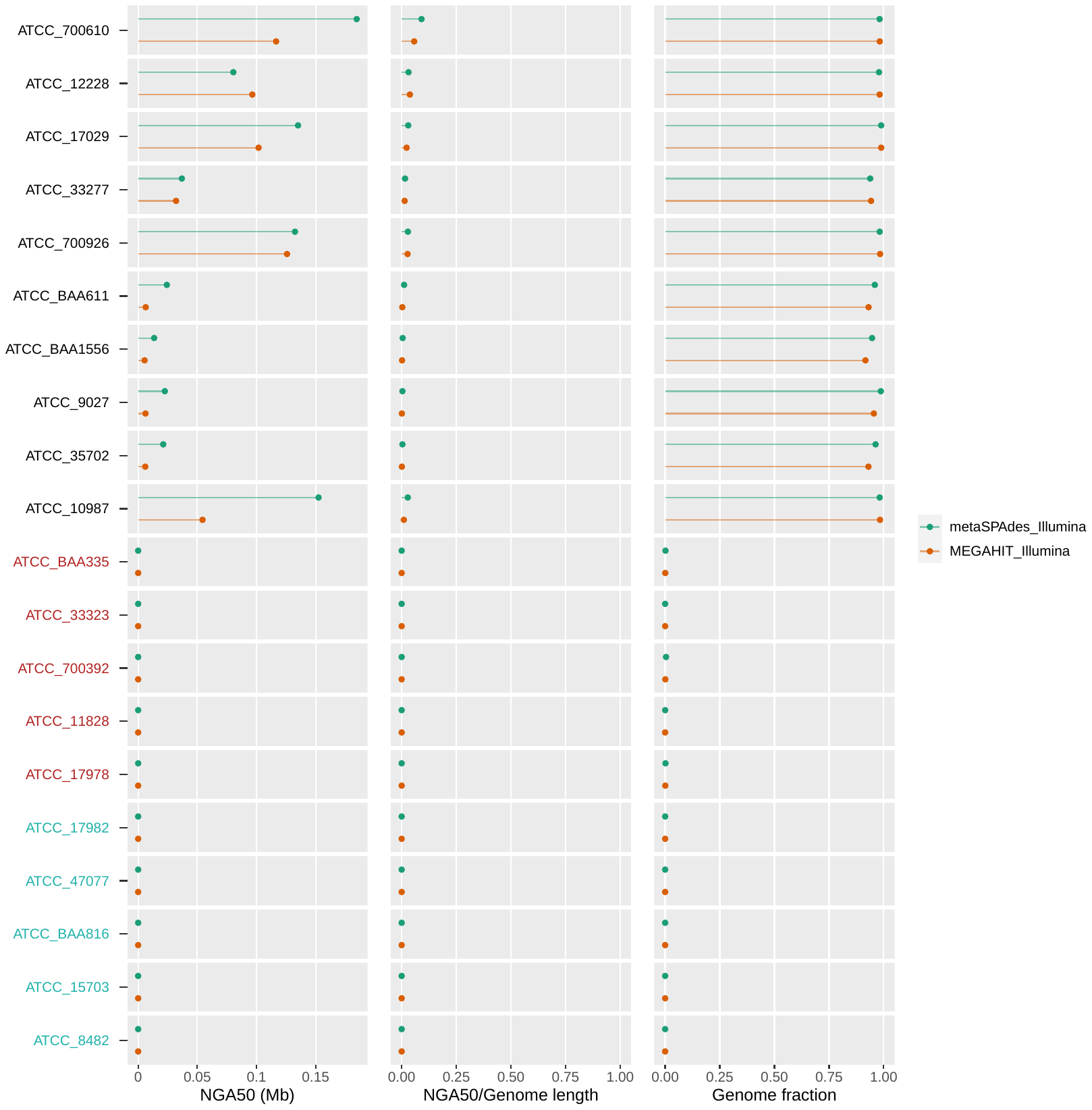

**Supplementary Figure 6.** Contig continuities and genome fractions (*GF*) for the short-read assemblies on the ATCC20 dataset. The red and green colors on the y-axis represent low- and ultra-low abundance species, respectively. The suffixes in the figure legend indicate the corresponding sequencing platforms.

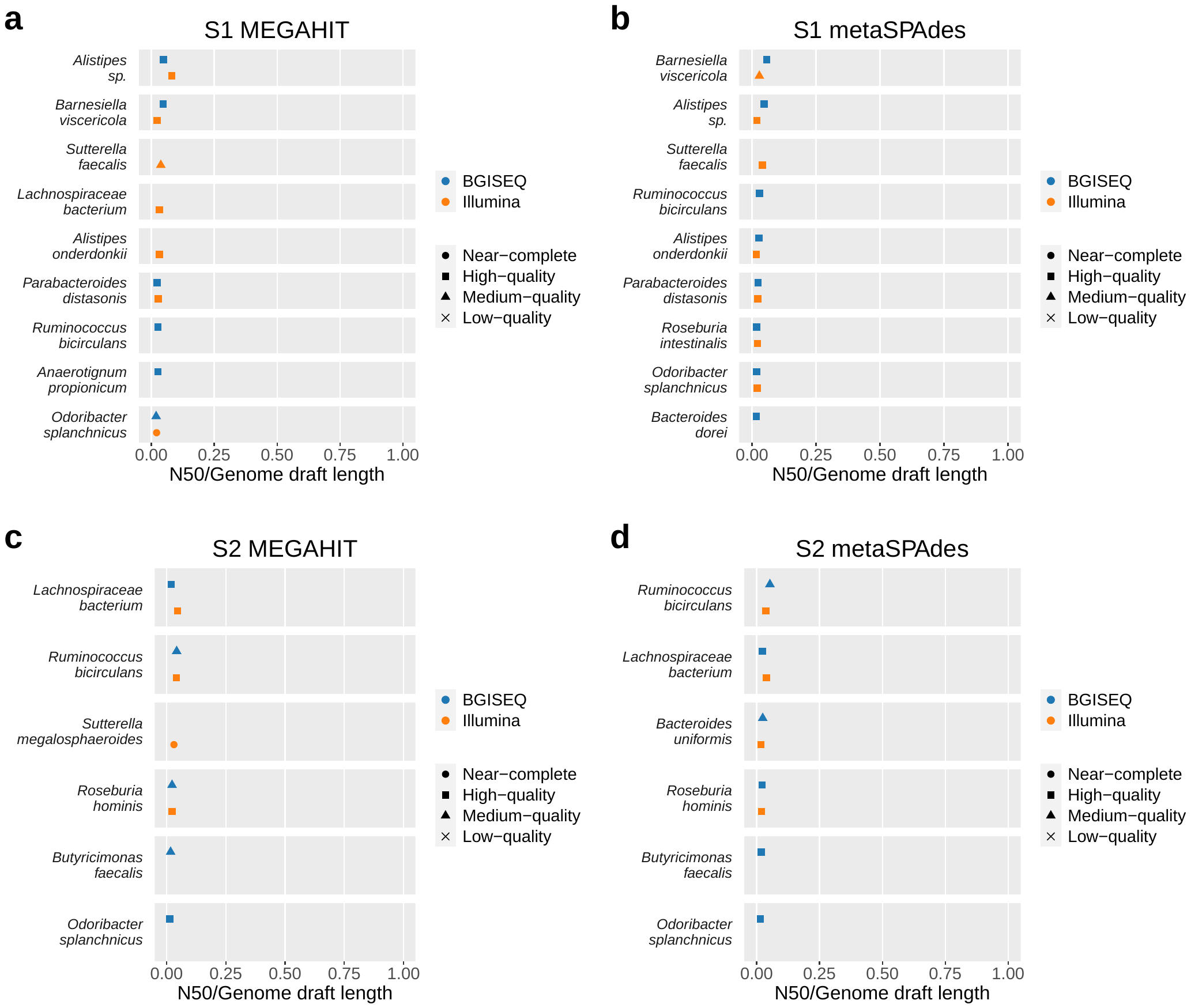

**Supplementary Figure 7.** The MAG annotations for S1 and S2 generated by BGISEQ-500 and Illumina HiSeq 2500.

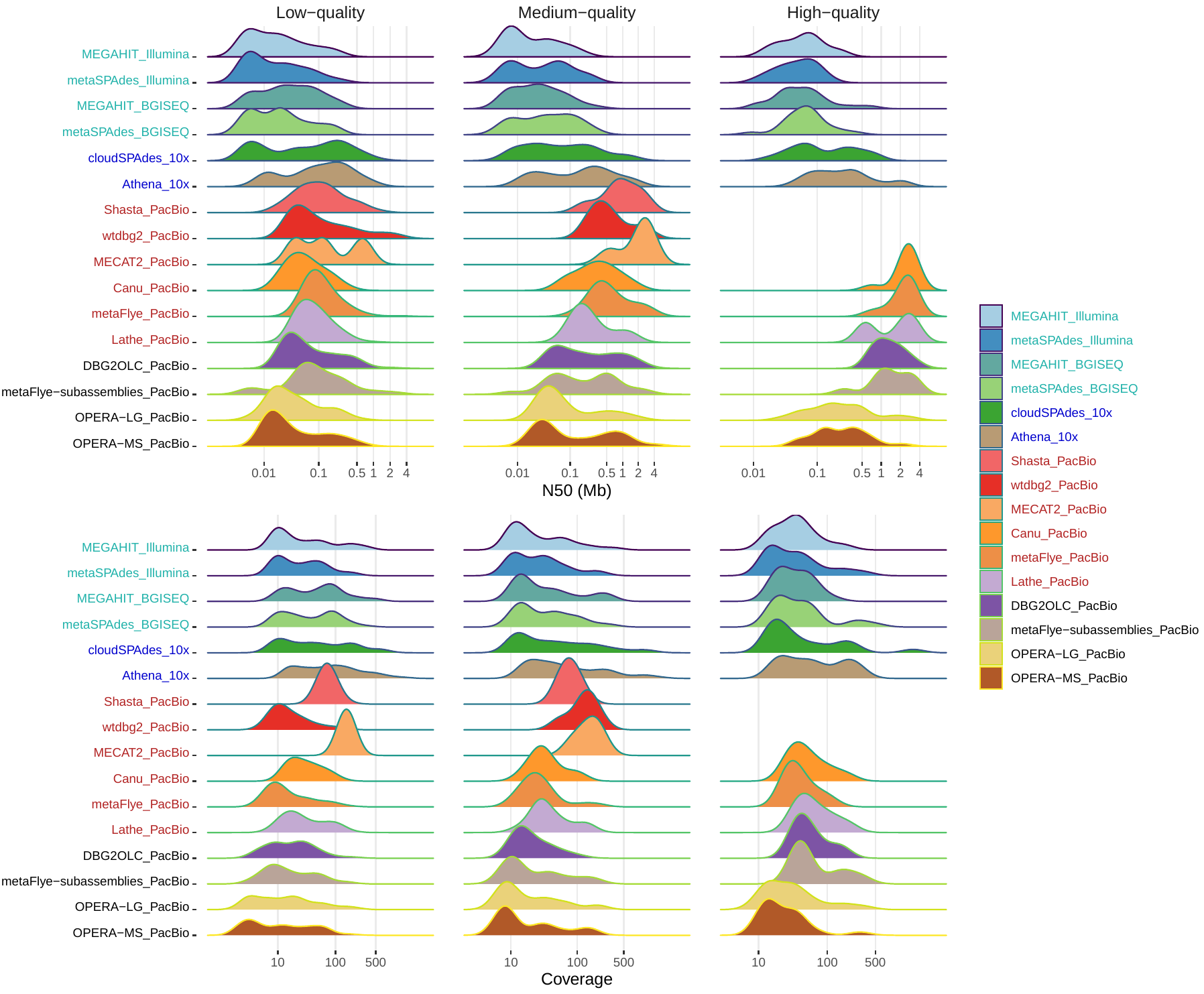

**Supplementary Figure 8.** MAG N50s and coverage distribution for S1. The suffixes in the figure legend indicate the corresponding sequencing platforms. The green, blue, red, and black colors denote short-read, linked-read, long-read, and hybrid assembly tools, respectively.

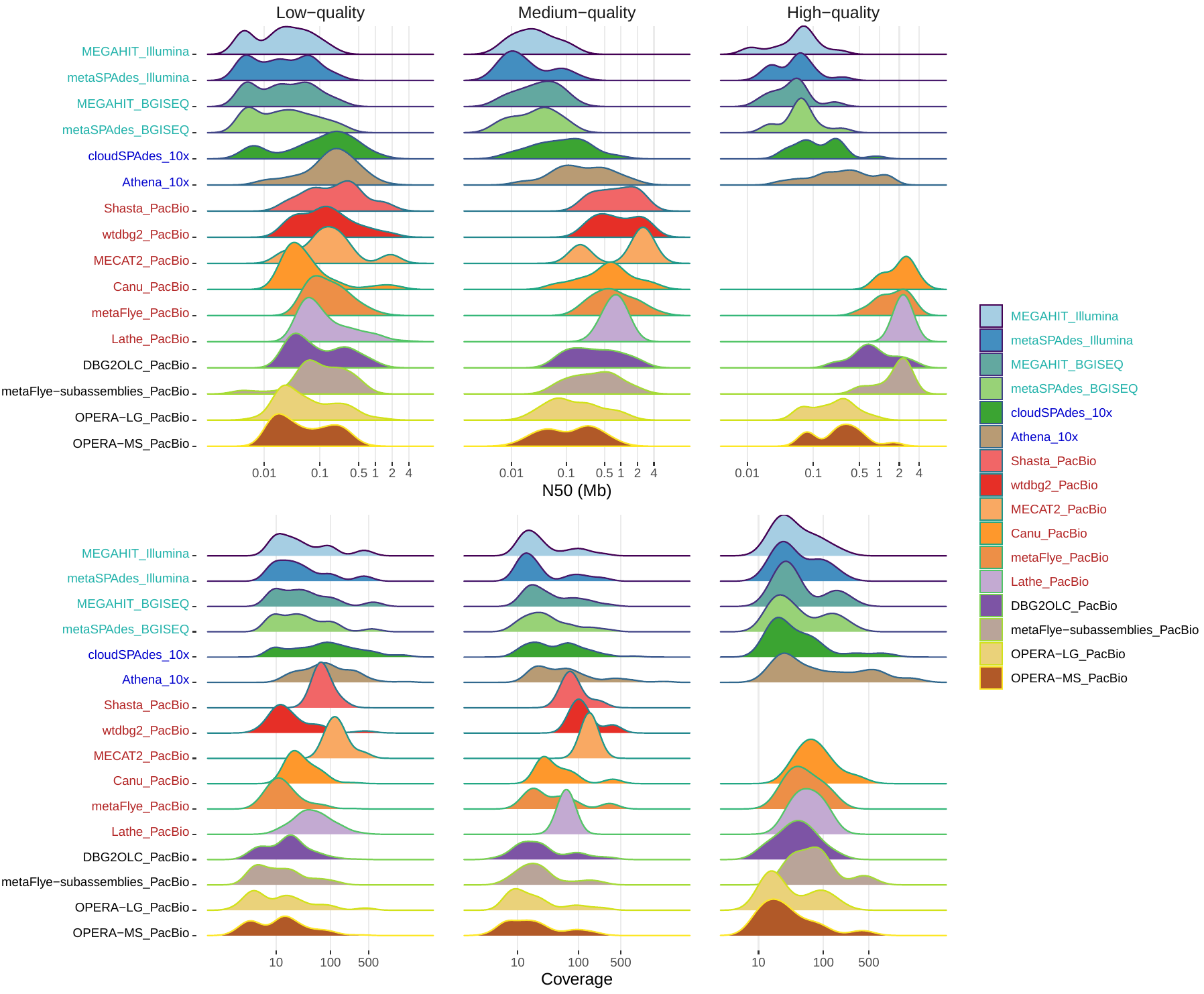

**Supplementary Figure 9.** MAG N50s and coverage distribution for S2. The suffixes in the figure legend indicate the corresponding sequencing platforms. The green, blue, red, and black colors denote short-read, linked-read, long-read, and hybrid assembly tools, respectively.

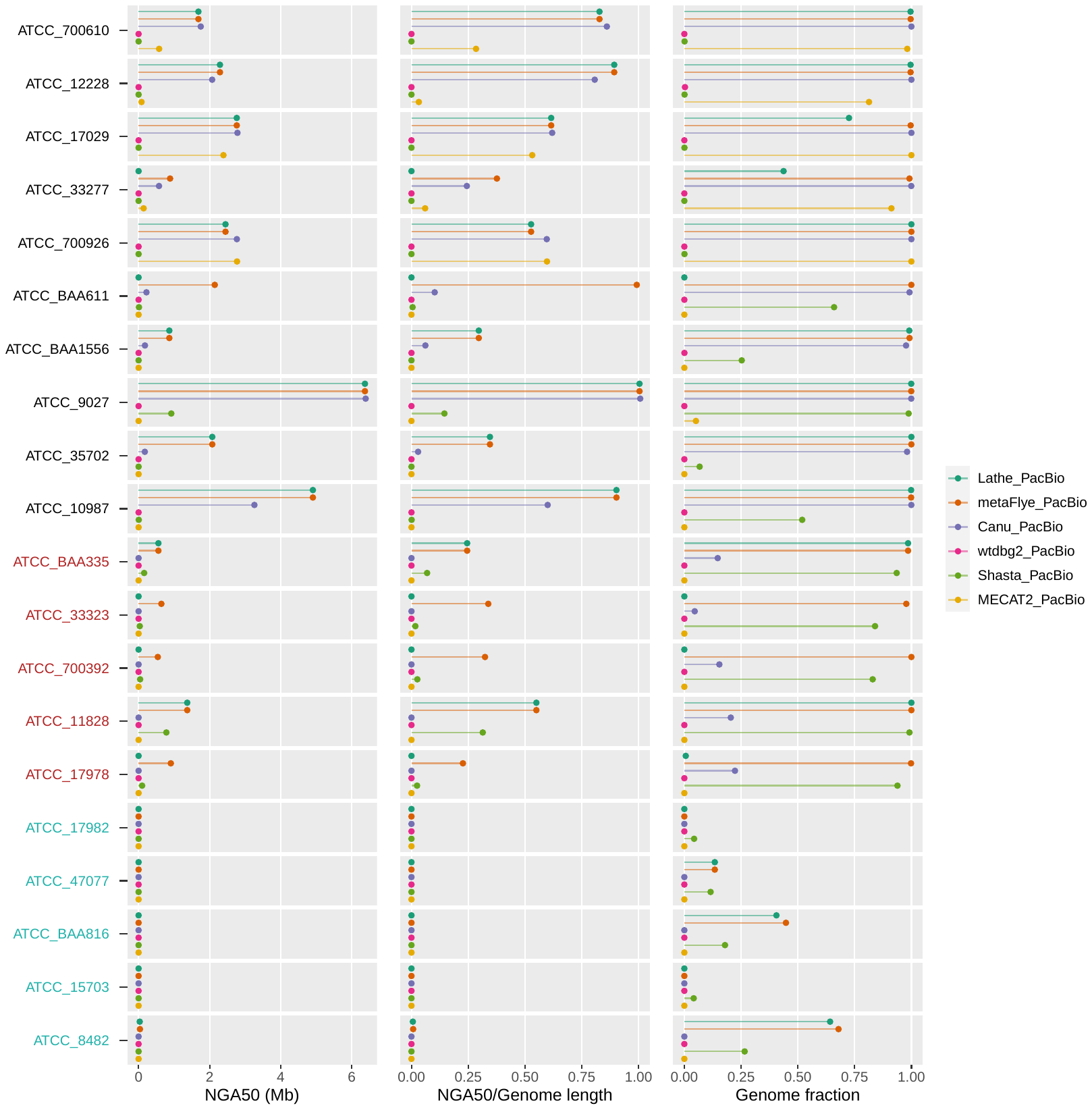

**Supplementary Figure 10.** Contig continuities and genome fractions (*GF*) for the long-read assemblies on the PacBio CLR dataset of ATCC20. The red and green colors on the y-axis represent low- and ultra-low abundance species, respectively. The suffixes in the figure legend indicate the corresponding sequencing platforms.

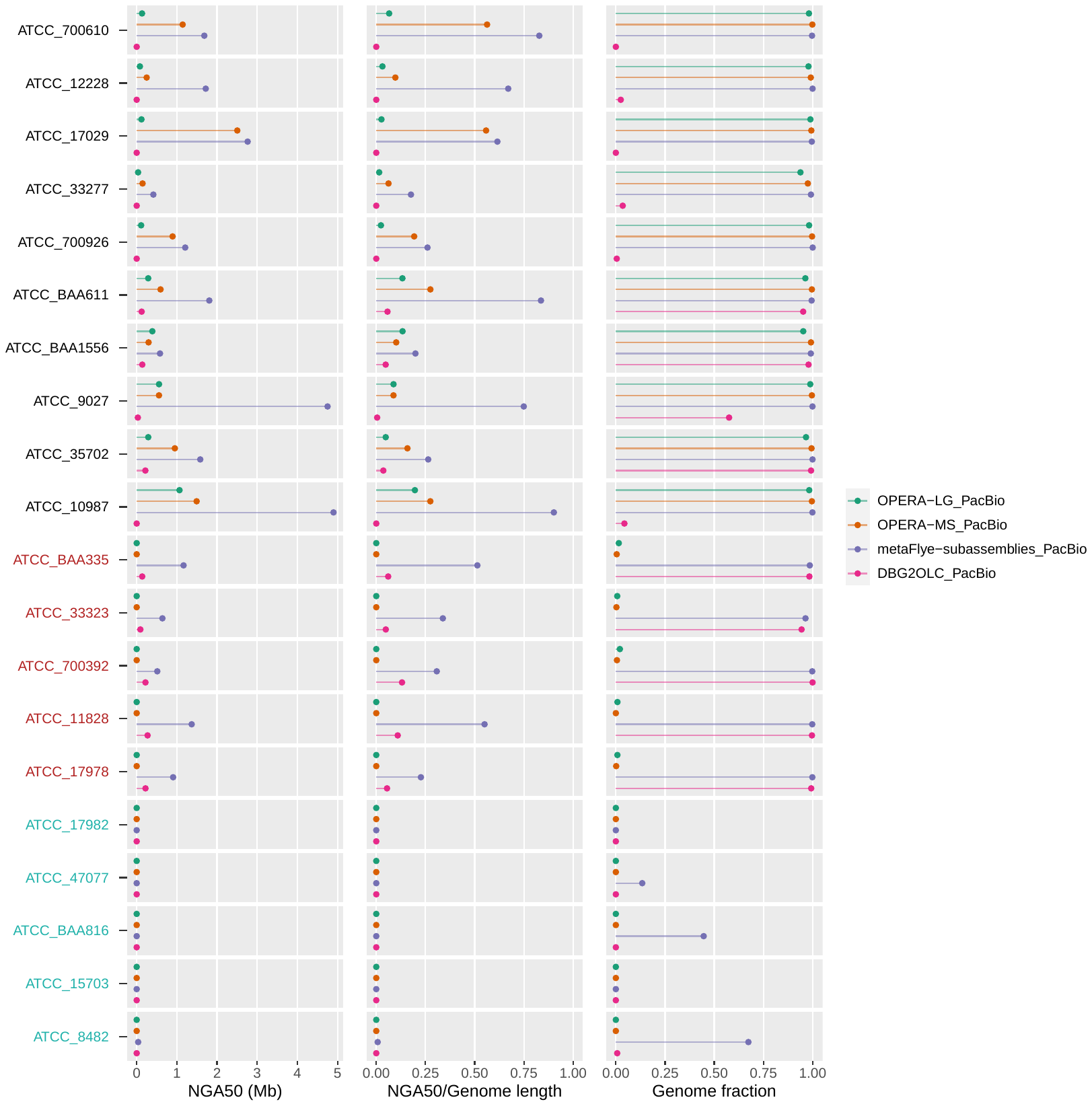

**Supplementary Figure 11.** Contig continuities and genome fractions (*GF*) for the hybrid assemblies on the ATCC20 dataset. The red and green colors on the y-axis represent low- and ultra-low abundance species, respectively. The suffixes in the figure legend indicate the corresponding sequencing platforms.

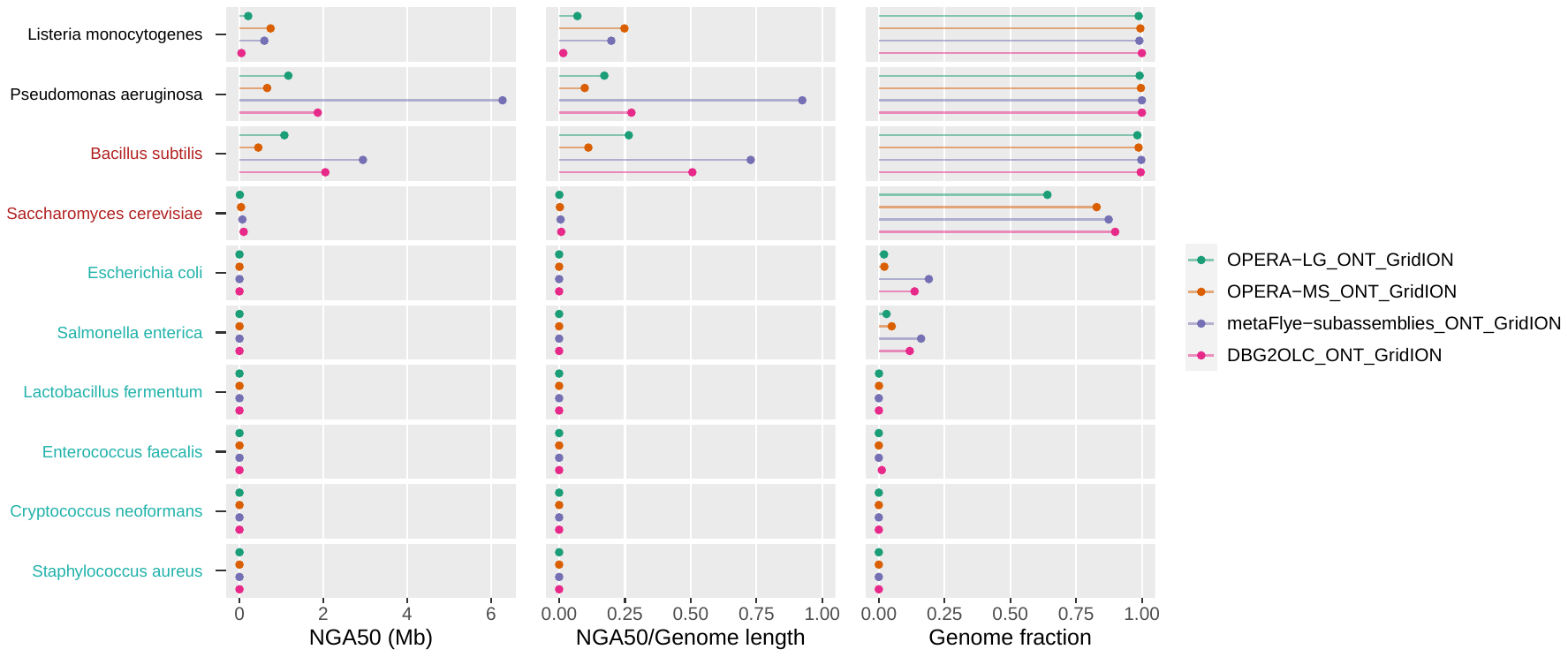

**Supplementary Figure 12.** Contig continuities and genome fractions (*GF*) for the hybrid assemblies (Illumina + ONT GridION) on the ZYMO dataset. The red and green colors on the y-axis represent low- and ultra-low abundance species, respectively. The suffixes in the figure legend indicate the corresponding sequencing platforms.

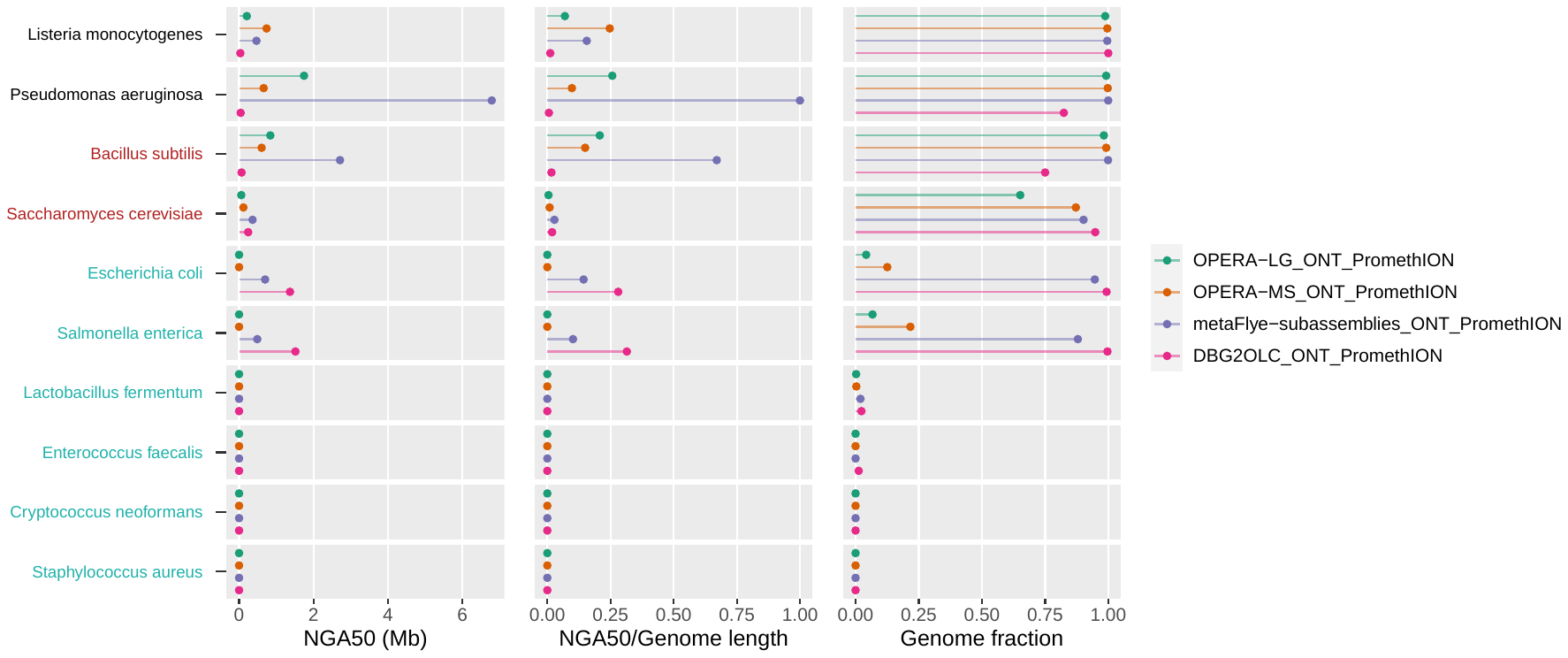

**Supplementary Figure 13.** Contig continuities and genome fractions (*GF*) for the hybrid assemblies (Illumina + ONT PromethION) on the ZYMO dataset. The red and green colors on the y-axis represent low- and ultra-low abundance species, respectively. The suffixes in the figure legend indicate the corresponding sequencing platforms.

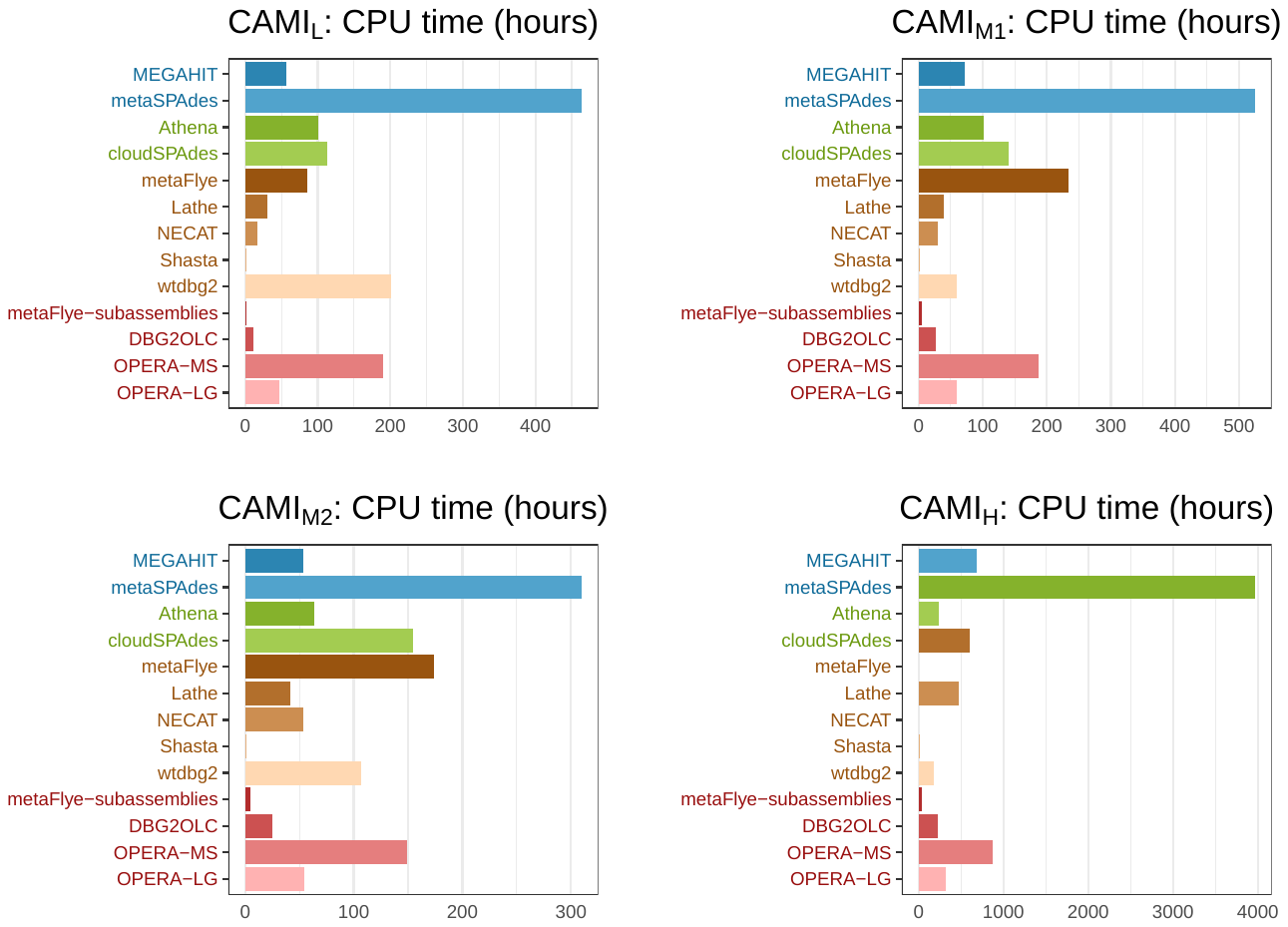

**Supplementary Figure 14.** CPU time consumed by the assembly tools in processing the CAMI datasets. We removed metaFlye and NECAT on CAMI_H_ because they exceeded the maximum memory limitation.

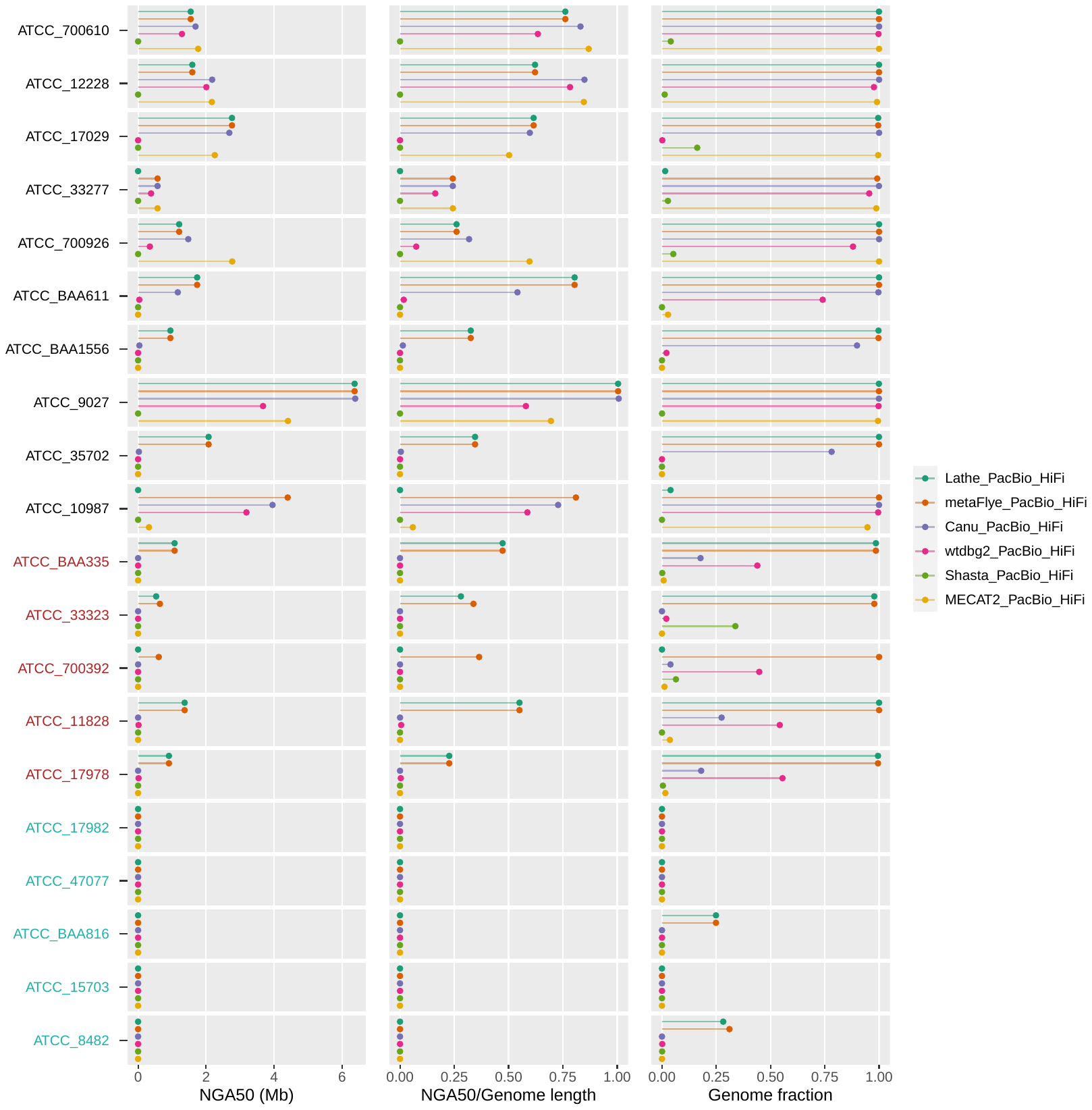

**Supplementary Figure 15.** Contig continuities and genome fractions (*GF*) for the long-read assemblies on the PacBio HiFi dataset of ATCC20. The red and green colors on the y-axis represent low- and ultra-low abundance species, respectively. The suffixes in the figure legend indicate the corresponding sequencing platforms.
